# Multiomic investigation of sugarcane mosaic virus resistance in sugarcane

**DOI:** 10.1101/2022.08.18.504288

**Authors:** Ricardo José Gonzaga Pimenta, Alexandre Hild Aono, Roberto Carlos Villavicencio Burbano, Marcel Fernando da Silva, Ivan Antônio dos Anjos, Marcos Guimarães de Andrade Landell, Marcos Cesar Gonçalves, Luciana Rossini Pinto, Anete Pereira de Souza

## Abstract

Sugarcane mosaic virus (SCMV) is the main etiological agent of sugarcane mosaic disease, which affects sugarcane, maize and other economically important grass species. Despite the extensive characterization of quantitative trait loci controlling resistance to SCMV in maize, the genetic basis of this trait is largely unexplored in sugarcane. Here, a genome-wide association study was performed and machine learning coupled to feature selection was used for the genomic prediction of resistance to SCMV in a diverse panel of sugarcane accessions. This ultimately led to the identification of nine single nucleotide polymorphisms (SNPs) explaining up to 29.9% of the phenotypic variance and a 73-SNP set that predicted resistance with high accuracy, precision, recall, and F1 scores. Both marker sets were validated in additional sugarcane genotypes, in which the SNPs explained up to 23.6% of the phenotypic variation and predicted resistance with a maximum accuracy of 69.1%. Synteny analyses showed that the gene responsible for the major SCMV resistance in maize is probably absent in sugarcane, explaining why such a major resistance source is thus far unknown in this crop. Lastly, using sugarcane RNA sequencing data, markers associated with the resistance to SCMV in sugarcane were annotated and a gene coexpression network was constructed to identify the predicted biological processes involved in SCMV resistance. This allowed the identification of candidate resistance genes and confirmed the involvement of stress responses, photosynthesis and regulation of transcription and translation in the resistance to this virus. These results provide a viable marker-assisted breeding approach for sugarcane and identify target genes for future molecular studies on resistance to SCMV.

## 1. Introduction

Sugarcane (*Saccharum* spp.) is highly economically important in tropical regions worldwide, as this plant is not only the world’s most important sugar-producing crop but also an important source of renewable energy obtained from its juice and bagasse (Carvalho-Netto *et al*., 2014; ISO, 2022). Brazil has been the leader of the global cultivation of sugarcane for many years and is currently responsible for approximately 40% of global production (FAO, 2022). However, sugarcane yield is threatened by several diseases, with sugarcane mosaic being of the most important at the global scale (Wu *et al*., 2012). In addition to the characteristic mosaic pattern displayed on the leaves, other symptoms of this disease include dwarfing, striping and streaking of culms, and shortening of internodes in highly susceptible genotypes (Gonçalves *et al*., 2007). In Brazil, this disease emerged in the beginning of the 20th century; it led to massive yield losses and drove the sugarcane industry to the brink of collapse in 1920-30. Damage caused by sugarcane mosaic disease has since been controlled with the employment of resistant cultivars and the adoption of several practices, such as the planting of healthy setts and roguing of nurseries and commercial fields. However, this disease is still a threat to sugarcane production, and resistance to it is a primary concern in breeding programs (Gonçalves *et al*., 2012).

Three viruses of the Potyviridae family are currently recognized as etiological agents of this disease in sugarcane: sugarcane mosaic virus (SCMV), sorghum mosaic virus, and sugarcane streak mosaic virus (Hall *et al*., 1998). SCMV, belonging to the *Potyvirus* genus, is a widespread species and the only one of these viruses found to naturally infect sugarcane in Brazil (Gonçalves *et al*., 2004, 2007, 2011). SCMV has been reported to cause sugarcane yield losses of up to 40-50% (Costa and Muller, 1982; Smith *et al*., 1992) while also reducing juice quality (Singh *et al*., 2003), sett germination and plant photosynthetic activity (Viswanathan and Balamuralikrishnan, 2005; Gonçalves *et al*., 2007). High yield losses arising from infection by this virus have led to the discontinuation of several sugarcane cultivars (Singh *et al*., 1997).

SCMV also infects many other closely related Poaceae species, including maize (*Zea mays*); this virus is responsible for extensive losses in maize yields, especially in Europe and China (Wu *et al*., 2012). As a result of being transmitted by various aphid species in a nonpersistent manner (Hassan *et al*., 2003), SCMV is very hard to control in the field, making host resistance an important resource to avoid damage caused by this virus. Thus, numerous quantitative trait locus (QTL) mapping studies have been performed to investigate the resistance of maize to SCMV; these studies resulted in the identification of two major and three minor QTLs controlling this trait in this species (Melchinger *et al*., 1998; Xia *et al*., 1999; Xu *et al*., 1999; Dußle *et al*., 2000, 2003; Zhang *et al*., 2003; Wu *et al*., 2007; Liu *et al*., 2009; Soldanova *et al*., 2012). Together, the major loci, named *Scmv1* and *Scmv2*, usually explain up to ∼60-70% of the phenotypic variance observed for resistance (Xia *et al*., 1999; Dußle *et al*., 2000; Soldanova *et al*., 2012). Recently, researchers have finely mapped the location of these QTLs and identified the causal genes responsible for resistance in maize (Ding *et al*., 2012; Tao *et al*., 2013; Li *et al*., 2016; Liu *et al*., 2017).

However, with respect to sugarcane, data on resistance to this virus are scarce. A few phenotypic studies have been performed in Brazil: specifically, researchers have evaluated the genotypic correlation of this disease incidence in sugarcane families (Xavier *et al*., 2013) and screened diverse genotypes for resistance (da Silva *et al*., 2015a, b). In addition, three marker– trait association studies have been carried out targeting SCMV resistance in this crop (Barnes *et al*., 1997; Pinto *et al*., 2013; Burbano *et al*., 2022); however, most included very few genotypes (≤50), and all employed dominantly scored markers. This apparent disparity between the information on SCMV resistance available for sugarcane and maize can be partially explained by the larger economic importance of the latter species in several countries; however, another important factor, i.e., sugarcane’s genomic complexity, has an effect. Modern cultivars are derived from a few crosses between two highly autopolyploid species, *Saccharum spontaneum* (2*n* = 5x = 40 to 16x = 128; *x* = 8) (Panje and Babu, 1960) and *Saccharum officinarum* (2*n* = 8x = 80; *x* = 10) (D’Hont *et al*., 1998). These hybrids have large (D’Hont *et al*., 1998), highly polyploid (D’Hont and Glaszmann, 2001), aneuploid (Sforça *et al*., 2019) and duplicated (Aono et al., 2021) genomes that hinder sugarcane breeding research. Additionally, studies suggest that the majority of sugarcane traits are controlled by many small-effect loci (Gouy *et al*., 2015; Fickett *et al*., 2019; Pimenta *et al*., 2021). However, given the existence of *Scmv1* and *Scmv2* in maize, it is odd that no major loci controlling SCMV resistance in sugarcane have been identified.

In view of this crop’s complex genome and the high impact of SCMV on its yield, the exploration of novel methodologies is required for the investigation of sugarcane’s resistance to this virus. This study aimed to identify markers associated with SCMV resistance and provide insights into its molecular basis through the use of state-of-the-art genomic and transcriptomic approaches. To achieve this, a panel of *Saccharum* accessions was assessed by phenotyping for SCMV resistance in the field and was genotyped via genotyping by sequencing (GBS), enabling the discovery of single nucleotide polymorphisms (SNPs) with information on allele proportion (AP) and position in a monoploid set of chromosomes of *S. spontaneum*. These data were used to perform a genome-wide association study (GWAS) to identify markers associated with SCMV resistance and to predict genotype attribution to resistant or susceptible groups by the use of machine learning (ML) coupled with feature selection (FS). Associated markers were genotyped on additional accessions previously assessed for SCMV resistance for validation and subsequently annotated by the use of a newly assembled sugarcane transcriptome. This allowed the incorporation of SCMV-associated genes into a coexpression network and thus a broader investigation of the molecular basis underlying sugarcane resistance to this virus.

## 2. Results

### 2.1. Panel phenotyping and genotyping

Ninety-seven sugarcane accessions were evaluated for the presence and severity of SCMV symptoms in two consecutive years. A skew in the distribution of the data toward the absence of symptoms was observed; despite normalization procedures, these data did not follow a normal distribution as indicated by a Shapiro–Wilk test (p = 2.2e-16). Based on the occurrence of SCMV symptoms, the panel could be divided into two groups: 62 resistant genotypes, which did not present symptoms in any block or year, and 35 susceptible genotypes, which presented symptoms on at least one occasion.

Following the construction and sequencing of a GBS library, 3,747,524, 3,152,409, and 569,360 biallelic SNPs were identified using FreeBayes, SAMtools and the TASSEL-4-POLY pipeline, respectively. After filtering procedures were performed and examining the intersection between tools, 37,001 of these markers were found to be called by TASSEL4-POLY and at least one of the other tools; thus, these markers constituted the final set of reliable SNPs.

### 2.2. Association analyses

#### 2.2.1. Mixed modeling

Data from 92 accessions of the panel were subjected to mixed modeling genome-wide association analyses on GWASpoly, and six different marker-effect models were used. Q-Q plots generated by these analyses can be found in Figure S2. In general, most models showed an appropriate profile of inflation of p values; exceptions disregarded for further analyses included the general model, which presented insufficient control of inflation, and the simplex dominant alternative model, which presented deflation. A stringent significance threshold (p < 0.05 corrected by the Bonferroni method) was used to establish 20 significant marker–trait associations, some of which were highly significant (Figure 1); the r^2^ values of associations ranged from 0.017 to 0.299 (Table S3). Several markers were associated with SCMV resistance according to more than one model, and nine nonredundant markers were representative of all associations.

**Figure 1.**
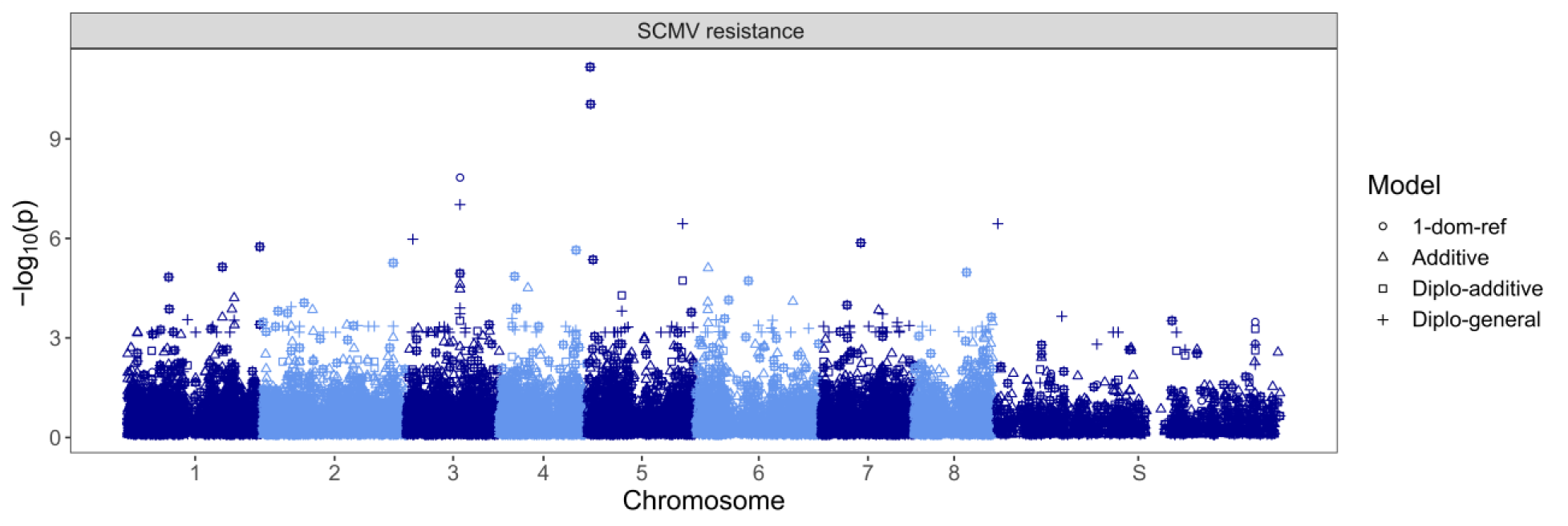
Manhattan plots generated from association analyses in which the best linear unbiased predictor (BLUP) values of SCMV symptom severity were used. Four different models were used: additive, simplex dominant reference (1-dom-ref), diploidized general (diplo-general) and diploidized additive (diplo-additive) models. On the x-axis, S represents scaffolds not associated with any of the *Saccharum spontaneum* chromosomes.

#### 2.2.2. ML coupled with FS

Eight ML algorithms for predicting the attribution of sugarcane genotypes to SCMV-resistant or SCMV-susceptible groups based on genotypic data were tested. When assessing their potential for this task when the full marker dataset was used, the predictive accuracies ranged from 52.8 (DT) to 66.9% (RF), with a mean of 60.3% (Table 1 and Figure S3). The remaining metrics evaluated showed much inferior results, with means of 21%, 26.8% and 20.7% found for precision, recall and F1 score, respectively. GP performed particularly poorly, with the mean of measurements equal to zero for these three metrics (Table 1 and Figures S4-S6).

**Table 1.** Predictive ability of machine learning (ML) models for predicting SCMV resistance before and after feature selection (FS). The ML models tested were adaptive boosting (AB), decision tree (DT), Gaussian naive Bayes (GNB), Gaussian process (GP), K-nearest neighbor (KNN), multilayer perceptron neural network (MLP), random forest (RF) and support vector machine (SVM).

| Model | Accuracy | Precision | Recall | F1 |
| --- | --- | --- | --- | --- |
| **Before Feature Selection** |  |  |  |  |
| **AB** | 60.7 | 23.8 | 36.6 | 27.3 |
| **DT** | 52.8 | 30.7 | 29.5 | 29.2 |
| **GNB** | 54.4 | 30.9 | 31.7 | 28.9 |
| **GP** | 66.3 | 0.00 | 0.00 | 0.00 |
| **KNN** | 59.6 | 18.1 | 30.9 | 21.6 |
| **MLP** | 55.3 | **54.6** | 38.3 | **43.0** |
| **RF** | **66.9** | 8.2 | **38.8** | 13.2 |
| **SVM** | 66.6 | 1.4 | 8.4 | 2.4 |
| **Mean** | 60.3 | 21.0 | 26.8 | 20.7 |
| **After Feature Selection** |  |  |  |  |
| **AB** | 86.2 | 70.0 | 87.9 | 76.5 |
| **DT** | 66.4 | 50.4 | 52.1 | 49.2 |
| **GNB** | 96.7 | 95.2 | 95.6 | 94.9 |
| **GP** | 93.2 | 81.7 | 98.2 | 88.4 |
| **KNN** | 95.7 | 87.3 | **100.0** | 92.8 |
| **MLP** | **99.7** | **100.0** | 99.3 | **99.6** |
| **RF** | 85.6 | 57.5 | 98.6 | 70.7 |
| **SVM** | 98.1 | 94.5 | **100.0** | 96.9 |
| **Mean** | 90.2 | 79.6 | 91.4 | 83.6 |

Therefore, three FS methods were used to reduce the marker dataset and improve model performance, and the SNPs identified by at least two of these methods were selected. This enabled the identification of a 73-SNP dataset, which led to considerable increases in all the metrics of all the models. With the reduced dataset, a mean accuracy of 90.2%, with a maximum of 99.7% using MLP, was obtained (Table 1 and Figure S3). Even more pronounced increases were observed for the other metrics: the mean precision was 79.6%, with a maximum of 100% with MLP (Table 1 and Figure S4); the mean recall was 91.4%, with a maximum of 100% with KNN and SVM (Table 1 and Figure S5); and the mean F1 score was 83.6%, with a maximum of 99.6 with MLP (Table 1 and Figure S6). ROC curves and their AUCs supported the promising results of FS in the predictive task. When all the markers were used, all the models presented ROC curves rather close to the level associated with chance alone, with AUCs ranging between 0.46 and 0.57 (Figure 2A). However, when markers selected by FS were used, most ROC curves indicated much better model performance, with AUCs of up to 0.99 (Figure 2B). Only DT and GP did not show appreciable increases in the AUC; thus, these models were excluded as appropriate methods for genomic prediction in this case.

**Figure 2.**
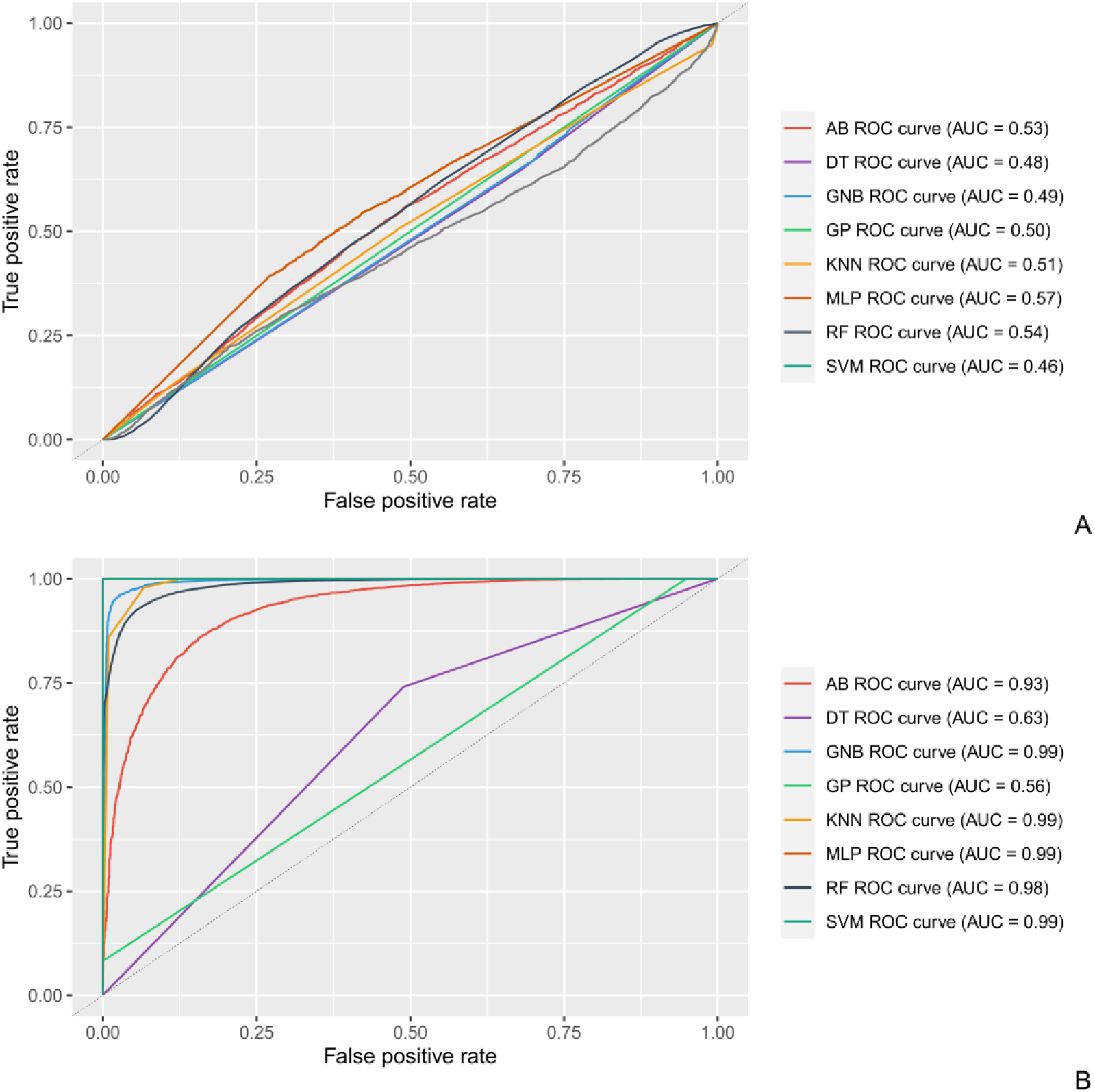
Receiver operating characteristic (ROC) curves and area under the curve (AUC) results concerning the performance of machine learning models for predicting SCMV resistance in which the full marker dataset (A) and markers selected by feature selection (FS) (B) were used. The machine learning models tested were adaptive boosting (AB), decision tree (DT), Gaussian naive Bayes (GNB), Gaussian process (GP), K-nearest neighbor (KNN), multilayer perceptron neural network (MLP), random forest (RF) and support vector machine (SVM).

### 2.3. Marker validation

Two groups of sugarcane genotypes previously assessed for SCMV resistance were genotyped via the MonsterPlex technology to validate markers identified in the association panel. The sequencing of the MonsterPlex library generated a total of 38,581,797 single-end reads, 99.8% of which presented a mean Q-value greater than 30; these values remained consistently high for the first 100 bases of the reads (Figure S7A). These data encompassed 81 of the 92 samples sent for analysis; DNA from genotypes 4, 5, 18, 68, 69, 70, 71, 73, 76, 77, and 79 (see Table S2) did not amplify well, and consequently, these samples were absent in the sequencing results. Interestingly, ten out of these samples were represented by *S. officinarum* F1 accessions, with only genotype 18 representing a hybrid variety. After trimming was performed, 38,574,693 reads were retained, 99.9% of which had a mean Q-value greater than 30 (Figure S7B). Using SAMtools and FreeBayes, 53 out of the 82 target SNPs (64.6%) could be called.

Seven of these SNPs were identified by GWAS as being significantly associated with SCMV resistance. When associations involving these markers were tested by linear models, the r^2^ values were overall lower than those of the original panel and were frequently close to zero, especially in the small wild accession panel. However, these values remained positive and reached as high as 0.236 (Table S4). The remaining 46 SNPs identified belonged to the reduced 73-SNP dataset identified by FS. These markers were applied to the eight ML models tested, which resulted in a mean accuracy of 61.6%, with a maximum of 69.1% by the RF model. This model was also among the top-ranking ones in terms of precision (68.6%), recall (94.1%) and F1 score (79.3%) for the identification of resistant genotypes (Table 2).

**Table 2.** Predictive accuracy, precision, recall and F1 scores of machine learning (ML) approaches employed to predict groups associated with SCMV resistance in the validation panel. The ML models tested were adaptive boosting (AB), decision tree (DT), Gaussian naive Bayes (GNB), Gaussian process (GP), K-nearest neighbor (KNN), multilayer perceptron neural network (MLP), random forest (RF) and support vector machine (SVM).

| **Model** | **Accuracy (%)** | **Precision (%)** | **Recall (%)** | **F1 (%)** |
| --- | --- | --- | --- | --- |
| **AB** | 61.7 | 64.7 | 86.3 | 73.9 |
| **DT** | 59.3 | 61.8 | 92.2 | 74.0 |
| **GNB** | 46.9 | 61.8 | 41.2 | 49.4 |
| **GP** | 65.4 | 65.3 | **96.1** | 77.8 |
| **KNN** | 64.2 | 63.8 | 100.0 | 77.9 |
| **MLP** | 63.0 | **69.1** | 74.5 | 71.7 |
| **RF** | **69.1** | 68.6 | 94.1 | **79.3** |
| **SVM** | 63.0 | 54.0 | 79.1 | 64.2 |
| **Mean** | 61.6 | 63.6 | 82.9 | 71.0 |

### 2.4. Synteny analyses

To assess the presence of the two major SCMV resistance QTLs from maize, *Scmv1* and *Scmv2*, in sugarcane, the CDSs of the causal genes at these loci were aligned against the *S. spontaneum* genomic sequence employed for SNP calling. Despite the close phylogenetic relation of the two species, no hits were found for *Scmv1*, indicating that this gene is likely absent from the *S. spontaneum* genome. Complementary searches in the genomes of an additional six sugarcane accessions also revealed no matches for this gene. The sequence of *Scmv2*, on the other hand, resulted in a 373-bp alignment with 88.7% identity and an E-value of 4.92e-127, corresponding to the Sspon.02G0027920-1A gene, which, like the causal gene at *Scmv2*, encodes an auxin-binding protein. This gene is located on chromosome 2A and is 1.7 kb away from the marker Chr2A_103190628, which was identified as being associated with SCMV resistance by FS (Figure 3).

**Figure 3.**
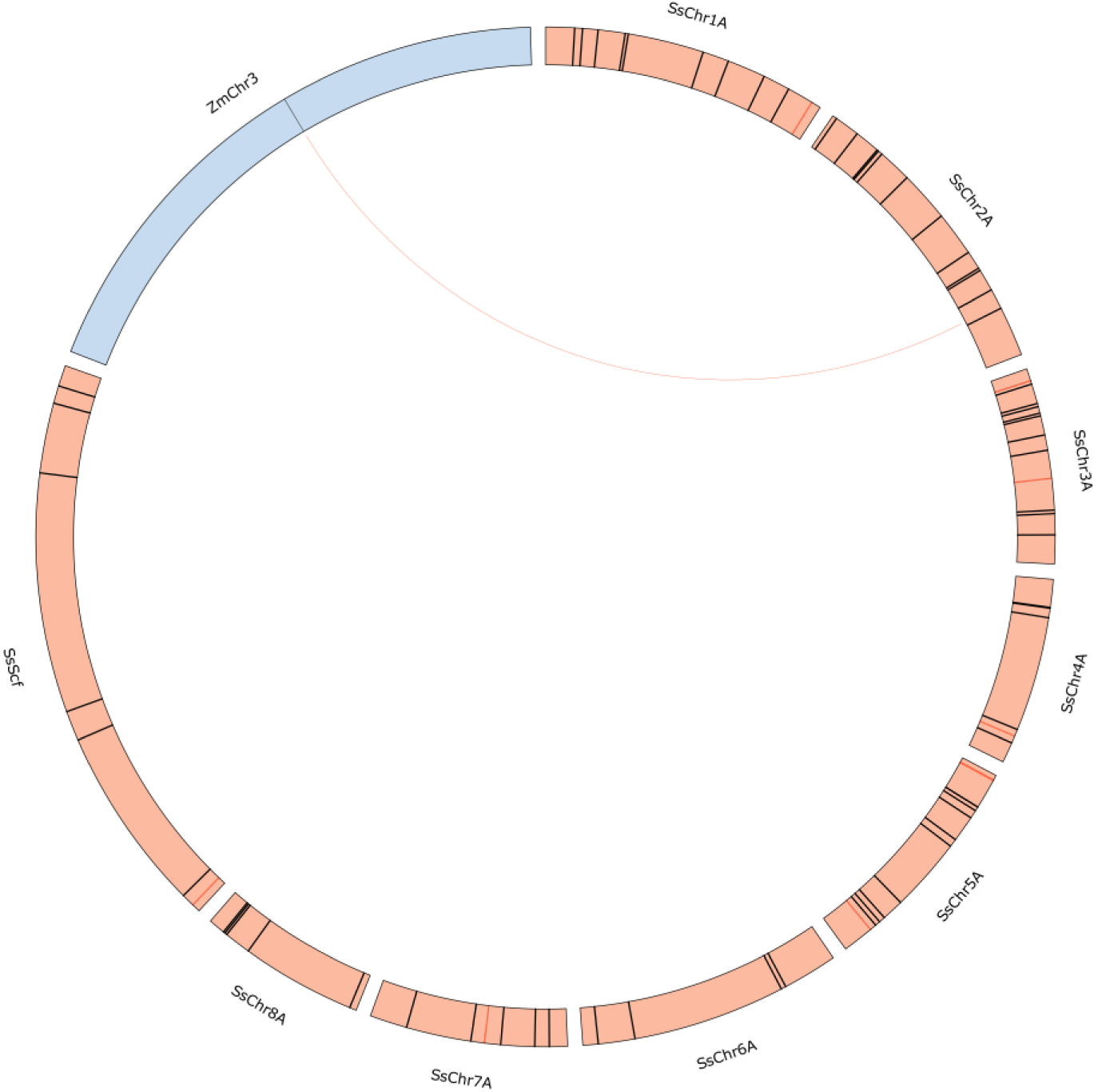
Synteny plot of *Scmv2* on chromosome 3 of *Zea mays* (blue) and *Saccharum spontaneum* A chromosomes (red). The red and black ticks represent markers associated with sugarcane mosaic virus (SCMV) resistance by association mapping and feature selection (FS), respectively.

### 2.5. Coexpression network construction and marker annotation

To assemble a de novo sugarcane transcriptome for marker annotation and expression analysis, more than two billion (2,477,287,294) sugarcane RNA sequencing (RNA-Seq) reads were retrieved from the SRA, 76% of which (1.9 billion) were retained after trimming. The transcriptomic reference assembled by Trinity comprised 611,480 transcripts with an N50 of 1,233 bp, represented by 212,076 longest isoforms (henceforth referred to as “genes”) with an N50 of 2,561 bp. The complete assembly contained 83.8% of conserved orthologs from green plants, as reported by BUSCO (Table S5). After quantification with Salmon, 131,615 genes were discarded for exhibiting very low expression. The remaining genes were used to construct a GWGCN. Using the UPGMA method, 64 functional modules were defined in this network, with sizes ranging from 58 to 32,980 genes and a mean size of 1,257.

To annotate the markers identified as associated with SCMV resistance through GWAS and FS, the transcriptome assembly was aligned against the *S. spontaneum* genome used for SNP calling, and the closest genes aligned upstream and downstream of each marker were retrieved. This enabled the association of 69 markers with 220 isoforms representing 84 genes. Thirty-five of these genes were located in 26 modules in the coexpression network; a summary of the alignment results is provided in Table S6. Among the annotated genes, a disease resistance protein associated with the Chr1A_90316612 marker was particularly interesting. To obtain better visualization of the biological processes associated with all the annotated genes, their GO-associated terms were retrieved and used for constructing a network using the REVIGO tool (Figure 4). The most prominent terms identified were linked to the regulation of transcription and translation, stress responses, and organismal development. The “modulation by virus of host process” term, which is associated with a peroxisomal oxidase and the marker Chr6A_86163774, was also displayed.

**Figure 4.**
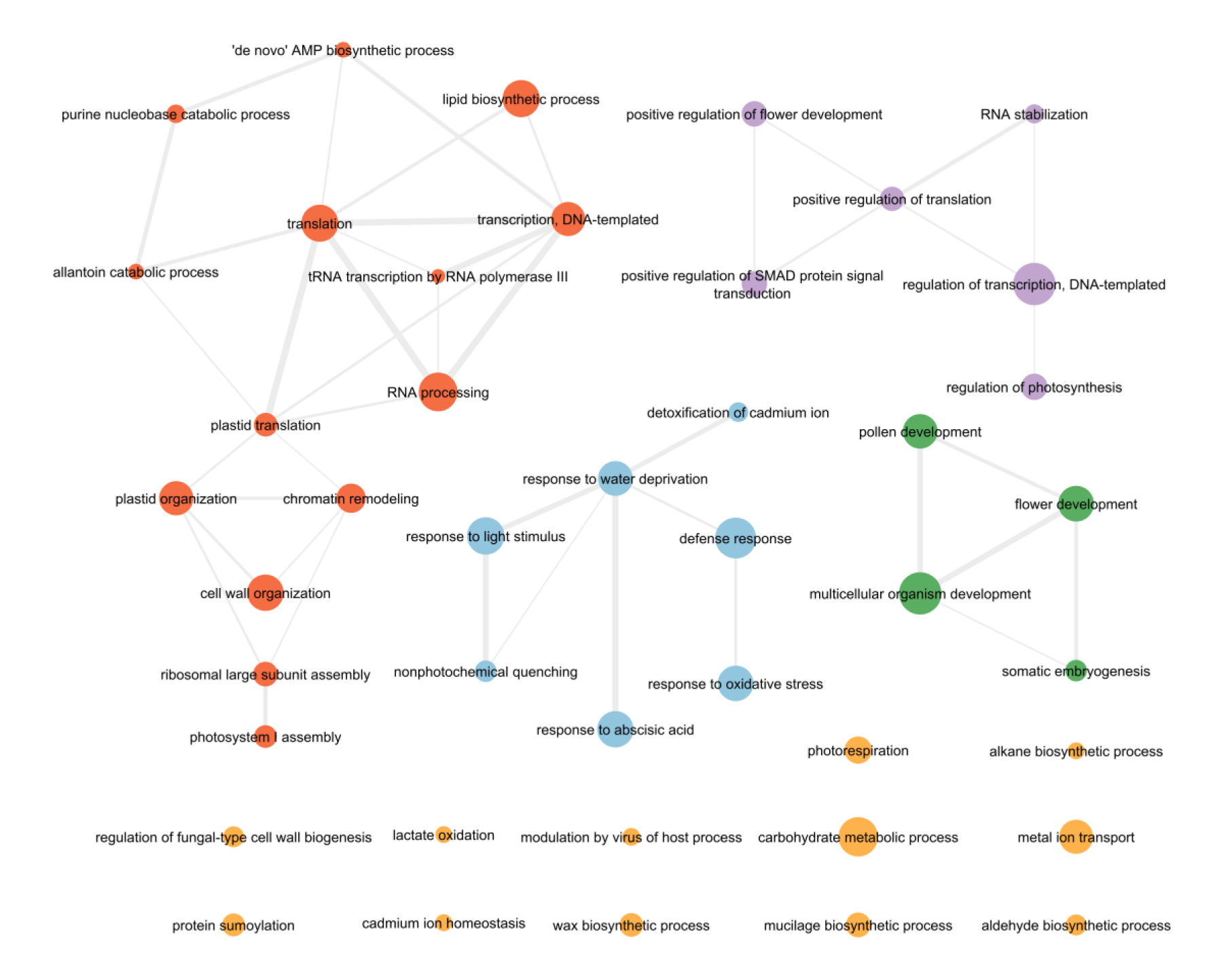
Network of Gene Ontology (GO) biological process terms obtained from genes associated with sugarcane mosaic virus (SCMV) resistance.

Additionally, the sequence of *Scmv2* from maize was aligned against the transcriptome, which resulted in several hits, the vast majority of which with TRINITY_DN5998_c0_g1. This gene was annotated as an auxin-binding protein and located within functional module 5 in the coexpression network—in which four genes close to SCMV resistance SNPs were also located. Among the hits with TRINITY_DN5998_c0_g1, the one with the highest E-value (1.07e-176) occurred in isoform 9, representing a 524-bp alignment with 88.1% identity. This and all isoforms of the gene also presented high-scoring alignments with the region containing Sspon.02G0027920-1A.

As a last strategy to investigate the biological processes involved in SCMV resistance, the GO terms of the 14,732 genes present in the 26 modules containing the genes associated with resistance were determined. Because these genes were initially associated with a very large number of GO terms (3,859 terms in the biological process category), a GO enrichment analysis using Fisher’s test with Bonferroni correction was performed before further procedures. The 117 terms resulting from this analysis were used to construct a TreeMap on REVIGO (Figure S8). The main terms found were distinct from those identified based only on the associated genes and included “response to salt stress”, “DNA integration”, “regulation of multicellular organismal process”, “photosynthesis”, and “seed germination”.

## 3. Discussion

Despite recent advances in sequencing technologies, genomic studies in sugarcane remain considerably hindered by the complexity of sugarcane’s genome (Thirugnanasambandam *et al*., 2018). Studies focusing on resistance to viruses are particularly limited due to sugarcane plants’ large size and vegetative propagation, which limit the size of controlled experiments and the number of genotypes that can be evaluated (da Silva *et al*., 2015; Pimenta *et al*., 2021). Moreover, the genotypes to be used in such assays can be made virus free only by tissue culture (Chatenet *et al*., 2001; Dewanti *et al*., 2016), a hard and time-consuming process. However, given the extensively reported economic impacts of SCMV infection on this crop’s yield (Costa and Muller, 1982; Bailey and Fox, 1987; Smith *et al*., 1992; Cronje *et al*., 1994; Singh *et al*., 2003), it is remarkable that this work represents the first genome-wide study targeting resistance to this pathogen in sugarcane.

Although our GWAS analysis did not reveal major loci controlling resistance to SCMV in sugarcane, it led to the identification of nine SNPs significantly associated with this trait, explaining a small (1.7%) to moderate (29.9%) percentage of the phenotypic variation. Compared to *Scmv1,* the major SCMV resistance QTL identified in maize, which explains 54-56% of the variation alone (Xia *et al*., 1999; Soldanova *et al*., 2012), these findings might seem modest. However, they fit in the upper range of those of other mapping studies in sugarcane; with the exceptions of resistance to brown and orange rust (Daugrois *et al*., 1996; Yang *et al*., 2018), markers explaining 10% or less of the phenotypic variation in traits of agronomic importance are common in this crop (Gouy *et al*., 2015; Fickett *et al*., 2019). Specifically, for resistance to SCMV, previous studies identified markers explaining 5-14% of the variance observed individually (Pinto *et al*., 2013; Burbano *et al*., 2022) or up to ∼40% together (Barnes *et al*., 1997). The apparent quantitative nature of resistance to SCMV in sugarcane attests to the limitations of traditional marker-assisted breeding in this crop and demonstrates the need for the deployment of high-throughput genotyping and specific methodologies for association analyses in sugarcane.

One finding from our study that contributes to the understanding of this panorama is the absence of a gene that is highly homologous to *ZmTrxh*, the causal gene at *Scmv1*, in the *S. spontaneum* genome. The expression of *ZmTrxh*, which encodes an atypical thioredoxin that acts as a molecular chaperone, is necessary to disrupt infection by SCMV (Liu *et al*., 2017). This phenomenon could involve SCMV’s RNA silencing suppressor helper-component proteinase (HC-Pro), which has been shown to interact with a maize ferredoxin (Cheng *et al*., 2008). These proteins are part of the ferredoxin–thioredoxin system, which is directly involved in photosynthesis (Buchanan, 1991) and might interfere with the suppression of silencing by SCMV HC-Pro and thus with resistance to this virus. Similar to other Poaceae (Liu *et al*., 2017), close orthologs of *ZmTrxh* are not present in *S. spontaneum*, resulting in the lack of this specific resistance mechanism in this crop. Since *ZmTrxh* is absent even in a few maize lines, its presence in other sugarcane genotypes cannot be completely ruled out. However, BLASTn alignments of its sequence were also performed against those of all other sugarcane genomes available to date, none of which returned significant alignments with the *ZmTrxh* sequence. Because sugarcane has a very narrow genetic basis (Panje and Babu, 1960), this gene is likely to be absent in other genotypes of commercial relevance.

In addition to performing a GWAS, ML algorithms coupled with FS were employed to predict genotype resistance or susceptibility to SCMV. Similar to previous works in which this genomic prediction methodology was applied to sugarcane to evaluate resistance to brown rust (Aono *et al*., 2020) and sugarcane yellow leaf virus (Pimenta *et al*., 2021), very promising results for several metrics were achieved. These results are considerably superior to those obtained by Barnes *et al*. (1997), who predicted sugarcane resistance to SCMV with an accuracy of 76% based on random amplified polymorphic DNA markers. Importantly, our results arose from a highly restricted SNP set obtained by FS, composed of only 73 markers, none of which had been identified by GWAS. A similar joint learning methodology that is based on FS and ML and combines classification and regression strategies has recently been shown to be highly suitable for the genomic prediction of several agronomic traits of sugarcane and polyploid forage grass species (Aono *et al*., 2022).

Unlike *Scmv1*, the causal gene at the second major SCMV resistance QTL from maize (*Scmv2*) has an ortholog in the *S. spontaneum* genome. Interestingly, one marker identified through FS (Chr2A_103190628) was found to be close (1.7 kb) to this region. Linkage disequilibrium is high in sugarcane, persisting for up to 2-3.5 Mb (Yang *et al*., 2019b; Pimenta *et al*., 2021). Thus, it is possible that this marker is linked to Sspon.02G0027920-1A, the *S. spontaneum* gene syntenic to the auxin-binding protein gene at *Scmv2* (Ding *et al*., 2012). This is an indication of the potential suitability of FS methodologies for the identification of QTLs, which is supported by other studies in which researchers analyzed traits controlled by many loci (Zhou *et al*., 2019).

To apply the findings of our study to sugarcane breeding, validation of the markers associated with SCMV resistance was performed. For sugarcane, validation of individual SNPs was successfully achieved only for resistance to orange rust (Yang *et al*., 2018; McCord *et al*., 2019). In the context of genomic prediction, SNP validation in sugarcane test populations was recently implemented by Hayes *et al*. (2021), who employed single-dose markers obtained through a SNP chip and achieved mean predictive accuracies of 29-47% for various agronomic traits. Due to the high cost of chip genotyping (De Donato *et al*., 2013; Bajgain *et al*., 2016) and the importance of including allele dose information in polyploid genetic studies (de Bem Oliveira *et al*., 2019; Aono *et al*., 2020), the MonsterPlex technology was chosen for the validation of our GBS-based markers. However, a few issues arose with this method, with failure in the amplification of a considerable percentage of sugarcane genotypes (∼22%) and loci (∼35%).

To some extent, this is expected from the technique, which does not guarantee the successful amplification and sequencing of all targets. The nonamplification of some genotypes might have been a consequence of the genomic reference used for SNP calling—the genome of *S. spontaneum*, a species different from that of almost all the genotypes for which amplification failed (*S. officinarum*). Because the majority of modern sugarcane cultivars that should be targeted by marker-assisted breeding are hybrids of these two species (Panje and Babu, 1960), failures in the amplification of whole genotypes are expected to be minimized. However, this highlights the importance of providing high-quality sequence data for genetically complex species such as sugarcane, which would certainly contribute positively to the research and breeding of this crop. Nevertheless, this low-cost targeted sequencing technology has the potential to be a viable approach for sugarcane marker-assisted breeding, especially if coupled to the ML-based genomic prediction approach used in this study, which effectively reduces the number of markers to be genotyped, contributing to the cost effectiveness of the process. The good results regarding predictive accuracy (69.1%), precision (68.6%), recall (94.1%) and F1 value (79.3%) obtained with the RF model are a strong indicator that this approach can be adopted for other traits of economic importance in sugarcane.

Another objective of the present work was to contribute to the elucidation of the molecular processes involved in sugarcane resistance to SCMV. To do so, RNA-Seq data was employed to annotate the markers identified as being associated with resistance and to construct coexpression networks to further investigate biological processes linked to them. Although RNA-Seq data from sugarcane plants infected with SCMV are available (Akbar *et al*., 2020), they include data from only two biological replicates, which equates to a very low sample number for network modeling—the WGCNA developers recommend a minimum of 15 samples to avoid noise and biologically meaningless inferences (Langfelder and Horvath, 2017). The summarization of GO terms from genes close to markers directly associated with SCMV resistance revealed a few general processes previously associated with responses to this virus; these included stress responses and the regulation of transcription and translation (Akbar *et al*., 2020; da Silva *et al*., 2020).

A more detailed examination of marker annotations revealed several genes previously linked to resistance to plant viruses; in many cases, these associations were established by RNA-Seq or proteomics. This is the case for allantoinases (Vuorinen *et al*., 2010), GLO oxidases (Varela *et al*., 2017), alpha-galactosidases (Naqvi *et al*., 2019), WD repeat-containing protein homolog genes (Şahin-Çevik *et al*., 2019), and pentatricopeptide repeat-containing proteins, which have also been associated with resistance to SCMV in maize through GWASs (Abdelkhalek *et al*., 2018; Gustafson *et al*., 2018). Similarly, Shen *et al*. (2021) identified a ribonuclease H protein gene at a QTL responsible for potyvirus resistance in soybean and showed that the expression of this gene was upregulated in resistant cultivars and influenced viral accumulation.

However, there is much more compelling evidence of associations with virus resistance in plants for other candidates identified in our study. For instance, resistance gene analogs with nucleotide-binding site leucine-rich repeat (NBS-LRR) motifs, such as RGA5, are often involved in disease resistance by their ability to recognize pathogenic effector proteins and induce effector-triggered immunity (Sekhwal *et al*., 2015). RGA5 specifically has been shown to bind effectors of a fungal pathogen in rice (Cesari *et al*., 2013), but NBS-LRR proteins also participate in the recognition and resistance to potyviruses (Ma *et al*., 2018; Xun *et al*., 2019). The existence of preliminary evidence of associations between polymorphisms in NBS-LRR protein genes and SCMV resistance in sugarcane (Brune and Rutherford, 2005) strengthens the hypothesis that RGA5 could in fact act as a resistance protein against infection by this virus.

Several genes that may represent susceptibility factors to SCMV were also annotated. For instance, a chloroplast carbonic anhydrase has been identified as a salicylic acid-binding protein that plays a role in the hypersensitive response of tobacco (Slaymaker *et al*., 2002). Furthermore, an *Arabidopsis* homolog of this protein was subsequently shown to interact with potyviral HC-Pro, weakening host defense responses and facilitating viral infection (Poque *et al*., 2018). A lower abundance of carbonic anhydrase was also associated with successful infection by *Tobamovirus* (Konakalla *et al*., 2021). In *Arabidopsis*, SCE1, a SUMO-conjugating enzyme, has been shown to interact with potyviral RNA-dependent RNA polymerase, and SCE1 knockdown resulted in increased resistance to turnip mosaic virus (Xiong and Wang, 2013). This protein also interacts with the replication initiator protein of begomoviruses and interferes with their replication (Castillo *et al*., 2004, 2007).

Additionally, three proteins that have chaperone activity and also participate in resistance to viruses were annotated. DNAJ and DNA-like proteins such as C76 and DNAJ 10 have been shown to interact with the coat protein of potyviruses, benefitting viral infection and replication (Hofius *et al*., 2007; Zong *et al*., 2020). Similarly, a heavy metal-associated isoprenylated plant protein was shown to interact with the *Pomovirus* movement protein, affecting virus long-distance movement (Cowan *et al*., 2018). Interestingly, transcripts of two chaperones and a heavy metal-associated isoprenylated protein differentially accumulated in response to SCMV in sugarcane (da Silva et al., 2020). Another protein annotated in the present study that has been shown to interact with the potyviral movement protein P3N-PIPO is a beta-glucosidase, possibly facilitating viral spread through the plant (Song *et al*., 2016); beta-glucosidase genes have also been found in QTLs for resistance to SCMV and other potyviruses (Gustafson *et al*., 2018; Rubio *et al*., 2019). Thus, it would be of great value to perform yeast two-hybrid assays including these host proteins and SCMV coat and movement proteins, the results of which could elucidate the involvement of these proteins in the replication and movement of SCMV.

Finally, the increase in the number of enriched GO terms associated with resistance through our GWGCN analysis sheds light on the complex network of biological processes involved in resistance to SCMV. The investigation of modules in coexpression networks can reveal sets of genes that are modulated together to execute specific functions; this is based on the “guilt-by-association” principle, which proposes that components (in our case, genes) with correlated biological functions tend to interact in networks such as GWGCNs (Oliver, 2000; Wolfe *et al*., 2005). According to the results of our analysis, biological processes enriched in SCMV resistance-associated modules included stress responses, regulation of transcription and translation, and a process that has long been known to be affected by SCMV infection (Irvine, 1971) but has not been featured by the analysis of genes directly associated with resistance— photosynthesis. Recent transcriptomic and proteomic studies have shown that infection by SCMV indeed affects the regulation of genes and proteins involved in these processes (Wu *et al*., 2013; Chen *et al*., 2017; Akbar *et al*., 2020; da Silva *et al*., 2020). Notably, the results of our coexpression network analyses indicate that the expression of genes identified in the present study as being associated with SCMV resistance are also associated with those controlling such processes; thus, those genes possibly play roles in their regulation during viral infection.

Our study indicates that resistance to SCMV in sugarcane has a more quantitative nature than in maize, which is in accordance with what has been observed for most traits in this crop. It also provides evidence that the ML-based strategy employed represents a viable approach for marker-assisted breeding in sugarcane; this strategy should therefore be assessed for its efficacy for other quantitative traits of economic importance. The annotation of identified markers via a transcriptomic assembly and analysis of gene coexpression networks showed that associated genes participate in key mechanisms of resistance to SCMV. These findings also revealed strong candidates for future investigation of resistance to this virus, which could help elucidate the molecular mechanisms involved in it.

## 4. Experimental procedures

### 4.1. Plant material

The plant material employed in the present study has been described elsewhere (Pimenta *et al*., 2021). The experimental population consisted of a panel of 97 sugarcane genotypes comprising wild accessions of *S. officinarum*, *S. spontaneum* and *Saccharum robustum*; traditional sugarcane and energy cane clones; and commercial cultivars from Brazilian breeding programs. The accession names and pedigree information are available in Table S1. A field experiment following a randomized complete block design with three blocks was established in May 2017 at the Advanced Center for Technological Research in Sugarcane Agribusiness located in Ribeirão Preto, São Paulo, Brazil (4°52’34” W, 21°12’50” S). Plants were grown in 1-meter-long three-row plots with row-to-row and interplot spacings of 1.5 and 2 meters, respectively. Each row contained two plants, totaling six plants of each genotype per plot. Infection by SCMV isolate RIB-2 (Burbano *et al*., 2022) was allowed to occur under natural conditions in conjunction with high inoculum pressure and a high incidence of aphid vectors.

### 4.2. Phenotyping

Plants were phenotyped in two cropping seasons: plant cane in February 2018 (9 months after planting) and ratoon cane in July 2019 (9 months after the first harvest). The severity of SCMV symptoms was assessed by 2-3 independent evaluators, who classified the top visible dewlap leaves in each plot by the use of a diagrammatic scale consisting of four levels of increasing intensity of mosaic symptoms (Figure S1).

The data normality was assessed by the Shapiro–Wilk test, and normalization was carried out using the bestNormalize package (Peterson, 2017) in R software (R Core Team, 2011). The best linear unbiased predictors (BLUPs) were estimated with the breedR R package (Munoz and Rodriguez, 2014) using a mixed model, as follows:

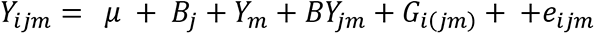

where *Yijm* is the phenotype of the i^th^ genotype considering the j^th^ block and the m^th^ year of phenotyping. The trait mean is represented by *μ*; fixed effects were modeled to estimate the contributions of the j^th^ block (*B_j_*), the m^th^ year (*Y_m_*) and the interaction between block and year (*BY_jm_*). Random effects included the genotype (*G*) and the residual error (*e*), representing nongenetic effects.

### 4.3. Genotyping

The library preparation and sequencing methods used were thoroughly described by Pimenta *et al*. (2021). Briefly, genomic DNA was extracted from the leaves and used for the construction of a GBS library following the protocol by Poland *et al*. (2012). For operational reasons, 94 out of the 97 genotypes of the panel were included in the library; genotypes 87, 88 and 95 were excluded (see Table S1). Two 150-bp single-end sequencing libraries were prepared, and their contents were sequenced on a NextSeq 500 instrument (Illumina, San Diego, USA). After checking the sequencing quality, three tools were used for SNP calling: SAMtools version 1.6 (Li *et al*., 2009), FreeBayes version 1.1.0-3 (Garrison and Marth, 2012) and the TASSEL4-POLY pipeline (Pereira *et al*., 2018). A monoploid chromosome set obtained from the *S. spontaneum* genome (Zhang *et al*., 2018) that included the A haplotype and unassembled scaffolds was used as a genomic reference. After variant calling, VCFtools version 0.1.13 (Danecek *et al*., 2011) was used to retain biallelic SNPs with a minor allele frequency of 0.1, a maximum of 25% missing data and a minimum sequencing depth of 50 reads. SNPs identified by TASSEL and at least one other tool were then selected, and the ratio between alleles (allele proportions, APs) was obtained for each marker.

### 4.4. Association analyses

#### 4.4.1. Mixed modeling

Association analyses were performed using mixed linear modeling in the GWASpoly R package (Rosyara *et al*., 2016). For these analyses, APs were transformed into genotypic classes with a fixed ploidy of 12 in the vcfR R package (Knaus and Grünwald, 2017), as proposed by Yang *et al*. (2019a). A realized relationship model (MM^T^) matrix (VanRaden, 2008), built in GWASpoly, was included as a random effect, and three principal components from a principal component analysis performed with genotypic data were included as fixed effects. Six marker-effect models were used for association analyses, namely, general, additive, simplex dominant reference, simplex dominant alternative, diploidized general and diploidized additive models. Q-Q plots of -log_10_(p) values of the markers were generated for all the models, and Manhattan plots were constructed for models with appropriate inflation profiles. The Bonferroni correction method with α = 0.05 was used to establish the significance threshold for associations. The phenotypic variance explained by each marker (r^2^) significantly associated with SCMV resistance was estimated using a linear model in R.

#### 4.4.2. ML coupled with FS

Following a genomic prediction approach previously employed for sugarcane (Aono *et al*., 2020; Pimenta *et al*., 2021), ML algorithms coupled with FS were used to predict the attribution of genotypes to two groups: those that presented mosaic symptoms at any block or year (susceptible) and those that did not present symptoms in any case (resistant). Eight ML algorithms implemented in the scikit-learn Python 3 module (Pedregosa *et al*., 2011) were tested: adaptive boosting (AB) (Freund and Schapire, 1997), decision tree (DT) (Quinlan, 1986), Gaussian naive Bayes (GNB) (Friedman *et al*., 1997), Gaussian process (GP) (Rasmussen, 2003), K-nearest neighbor (KNN) (Cover and Hart, 1967), multilayer perceptron (MLP) neural network (Popescu *et al*., 2009), random forest (RF) (Breiman, 2001) and support vector machine (SVM) (Cristianini and Shawe-Taylor, 2000). Three FS techniques were employed to obtain feature importance and create subsets of marker data: gradient tree boosting (FS1) (Chen and Guestrin, 2016), L1-based FS through a linear support vector classification system (FS2) (Cristianini and Shawe-Taylor, 2000) and univariate FS using analysis of variance (FS3) (Geurts *et al*., 2006), which were also implemented in scikit-learn. The markers selected by at least two of these FS methods were identified and used with the referred ML algorithms to classify genotypes as resistant or susceptible. To implement a cross-validation strategy, a stratified K-fold (k=5) repeated 100 times for different data configurations was used. The following metrics were evaluated: accuracy (proportion of correctly classified items), recall (items correctly classified as positive among the total quantity of positives), precision (items correctly classified as positive among the total items identified as positive), and the F1 score (the harmonic mean of precision and recall). The area under the receiver operating characteristic (ROC) curve (AUC) was also calculated for all the models using scikit-learn and plotted with the ggplot2 R package (Wickham, 2011).

### 4.5. Marker validation

Markers significantly associated with SCMV resistance were subjected to validation in two additional panels with sugarcane genotypes previously assessed for this trait. The first panel comprised 28 wild accessions, including representatives of *S. officinarum*, *S. spontaneum*, *S. robustum*, *Saccharum barberi*, and interspecific hybrids (da Silva *et al*., 2015a), and the second panel comprised 64 Brazilian varieties and elite clones from the three main sugarcane breeding programs in Brazil (da Silva *et al*., 2015b). These 92 genotypes (Table S2) were used for validation using MonsterPlex Technology (Floodlight Genomics, Knoxville, USA). DNA was extracted from leaves following the methods described by Aljanabi et al. (1999) or using the GenElute Plant Genomic DNA Miniprep Kit (Sigma–Aldrich, St. Louis, USA). DNA samples and marker flanking sequences were sent to Floodlight Genomics, where multiplex PCR was used to amplify ∼100-bp fragments containing markers, which were then sequenced on a HiSeq platform (Illumina, San Diego, USA). Trimmomatic version 0.39 (Bolger *et al*., 2014) was used to trim the single-end sequencing reads using a 5-bp sliding window with a minimum average Phred quality score of 20 and removing reads shorter than 30 bp. The trimmed reads were aligned to reference flanking sequences using Bowtie2 version 2.2.5 (Langmead and Salzberg, 2012), and SNP calling was performed using SAMtools and FreeBayes. After APs/genotypic classes were obtained for each locus, linear models in R were used to estimate marker r^2^ values for each panel, and ML models were used to predict resistance phenotypes as previously described.

### 4.6. Synteny analyses

For synteny analyses, the coding DNA sequences (CDSs) of the causal genes at *Scmv1* and *Scmv2* were retrieved from the MaizeGDB database (Portwood *et al*., 2019) and aligned against the *S. spontaneum* genome sequence using BLASTn (Altschul *et al*., 1990). Synteny plots were constructed using Circos software version 0.69.9 (Krzywinski *et al*., 2009). The *Scmv1* CDS was also aligned to the genome sequences of *S. spontaneum* Np-X (Zhang et al., 2022), *S. officinarum* LA Purple (SRA Bioproject accession PRJNA744175), and the hybrids SP70-1143 (Grativol *et al*., 2014), R570 (Garsmeur *et al*., 2018), SP80-3280 (Souza *et al*., 2019), and CC01-1940 (Trujillo-Montenegro *et al*., 2021).

### 4.7. Coexpression network construction and marker annotation

To annotate markers associated with SCMV and investigate their expression profile, RNA-Seq data supplied by Marquardt *et al*. (2019) was used. This study provided data from samples with five biological replicates, each made up of four to five bulked leaves, which were considered suitable for the construction of a highly robust coexpression network. Sequencing data were downloaded from the Sequence Read Archive (SRA; BioProject PRJNA474042) and trimmed with Trimmomatic version 0.39 (Bolger *et al*., 2014), with the default parameters.

A de novo transcriptome was assembled using Trinity version 2.5.1 (Grabherr *et al*., 2011), with the minimum contig length set to 300 bp. The completeness of the assembly was evaluated with BUSCO version 5.1.2 (Simão *et al*., 2015) using datasets of conserved orthologs from Viridiplantae. Annotations were performed with Trinotate (Bryant *et al*., 2017) and included homology searches of sequences in the UniProt database, domain identification according to information in the Pfam database, and predictions of signal peptides with SignalP and transmembrane domains using TMHMM. Salmon version 1.1.0 software (Patro *et al*., 2017) was used for transcript quantification, with the default parameters used. Genes with a mean of less than 5 transcripts per million (TPM) in at least one sample type were filtered out to avoid genes expressed at low levels, and genes with no variance across quantifications were excluded using the WGCNA package (Langfelder and Horvath, 2008).

A global weighted gene coexpression network (GWGCN) was constructed with WGCNA. Pairwise Pearson correlations of TPM values that considered a power function to fit a scale-free independence were used. For that, a soft threshold power beta estimation of 25, corresponding to an r² value of 0.85, was estimated and generated a scale-free topology model. Functional modules in the network were defined by the use of the unweighted pair group method with arithmetic mean (UPGMA) based on a topological overlap matrix and dynamic dendrogram pruning based on the dendrogram only.

To annotate markers associated with SCMV resistance and locate them in the network, the de novo transcriptome assembly was aligned against the *S. spontaneum* genomic reference used for SNP calling via BLASTn, and the closest genes upstream and downstream of each marker at a maximum distance of 2 Mb were retrieved. The following parameters were used: a minimum of 90% identity, a minimum E-value of 1e-50, and best hit algorithm overhang and edge values of 0.1. Similarly, the CDSs of the causal genes at *Scmv1* and *Scmv2* were aligned against the transcriptome assembly using BLASTn with the default parameters.

All genes present in the network modules containing genes associated with SCMV resistance were recovered and used for a Gene Ontology (GO) enrichment analysis with the topGO R package (Alexa and Rahnenfuhrer, 2010) in conjunction with Fisher’s test with a Bonferroni correction with α = 0.01. The REVIGO tool (Supek *et al*., 2011) was used for the visualization and analysis of GO categories of the genes associated with SCMV resistance and in enriched categories associated with the genes in the network modules.

## Acknowledgments

We thank Aline C. L. Moraes for assistance in constructing and sequencing the GBS library and Maicon Volpin for assistance with fieldwork.

## Author contributions

MCG, LRP and APS conceived the project and designed the experiments. RJGP, RCVB, MFS, IAA and LRP performed the phenotyping. RJGP and AHA performed the genotyping, analyzed the data and interpreted the results. RJGP wrote the manuscript. All the authors read and approved the manuscript.

## Data statement

All datasets analyzed during the current study are available online and referenced in the corresponding papers.

