## Supplementary Material for "Multiomic investigation of sugarcane mosaic virus resistance in sugarcane"

**Supplementary Tables**

**Table S1. List of genotypes in the association panel and their respective parents.**

| ID Number | Genotype | Female Parent | Male Parent |
|---|---|---|---|
| 1 | US571415 | *S. spontaneum* | SP701143 |
| 2 | Cana alho | *S. officinarum* | Unknown |
| 3 | IACSP012410 | R570 | IACSP933046 |
| 4 | IACSP046007 | PO8862 | IAC912195 |
| 5 | IACSP046059 | IACSP953018 | SP775181 |
| 6 | IACSP953028 | IACSP953028 | Unknown |
| 7 | NG5712 | *S. robustum* | Unknown |
| 8 | IACSP042516 | IAC912288 | CT961414 |
| 9 | Krakatau | *S. spontaneum* | Unknown |
| 10 | IN8482 | *S. spontaneum* | SP823530 |
| 11 | IACSP042508 | SP913059 | Unknown |
| 12 | IACSP955037 | SP842066 | IACSP973391 |
| 13 | IACCTC059630 | IACSP972053 | SP80185 |
| 14 | IACSP046053 | IACSP956114 | IAC873396 |
| 15 | IACSP993009 | CP701133 | Unknown |
| 16 | IS76155 | *S. officinarum* | RB855113 |
| 17 | IACSP012421 | RB835486 | SP847017 |
| 18 | IACSP042521 | IAC912288 | SP88721 |
| 19 | IACSP046058 | IACSP953018 | SP87365 |
| 20 | IACSP991020 | SP851682 | Unknown |
| 21 | IACSP045065 | IACSP956114 | RB72454 |

| | | | |
|---|---|---|---|
| 22 | IACCTC069767 | IACSP963069 | SP803280 |
| 23 | IACSP985046 | SP813251 | Unknown |
| 24 | IACSP018034 | IAC913093 | Unknown |
| 25 | IACSP993357 | SP921584 | Unknown |
| 26 | IACCTC059578 | White transparent | SP963092 |
| 27 | IACCTC055580 | CTC4 | IACSP933046 |
| 28 | IACSP976628 | RB835486 | SP931124 |
| 29 | IACSP976680 | SP842066 | Unknown |
| 30 | IACCTC059534 | IACSP953028 | Unknown |
| 31 | IACSP972084 | RB855035 | SP775181 |
| 32 | CTC15 | SP842025 | Unknown |
| 33 | IAC914168 | SP813137 | IAC873396 |
| 34 | IACBIO232 | RB855465 | CTC9 |
| 35 | IACBIO241 | RB855465 | SP963092 |
| 36 | IACBIO257 | NG26011 | IAC863154 |
| 37 | IACBIO266 | NG26011 | RB855156 |
| 38 | IACBIO270 | SES069 | Unknown |
| 39 | IACBIO271 | SES069 | SP80185 |
| 40 | IACBIO273 | SES069 | Unknown |
| 41 | IACBIO275 | SES069 | Unknown |
| 42 | IACBIO277 | SES069 | IACSP972077 |
| 43 | IACBIO279 | SES069 | IAC873396 |
| 44 | IACCTC053616 | IAC915155 | RB85453 |
| 45 | IACCTC056518 | SP911049 | SP832847 |
| 46 | IACCTC059552 | IACSP955011 | SP775158 |
| 47 | IACCTC059607 | IACSP972053 | SP963092 |

| | | | |
|---|---|---|---|
| 48 | IACCTC059634 | IACSP972053 | IACSP953050 |
| 49 | IACCTC061050 | CT943165 | IACSP952078 |
| 50 | IACCTC069708 | IACSP962042 | SP823697 |
| 51 | IACCTC069713 | IACSP955000 | SP87432 |
| 52 | IACCTC069741 | IACSP955000 | Unknown |
| 53 | IACSP012417 | RB835486 | RB835486 |
| 54 | IACSP012430 | IAC912195 | CT961414 |
| 55 | IACSP015501 | SP913059 | SP80185 |
| 56 | IACSP015519 | RB835486 | Unknown |
| 57 | IACSP018046 | R570 | Glagah |
| 58 | IACSP018082 | IAC913093 | Glagah |
| 59 | IACSP018158 | RB835486 | Unknown |
| 60 | IACSP022067 | SP973060 | Unknown |
| 61 | IACSP022125 | SP901161 | Unknown |
| 62 | IACSP023025 | SP924230 | Unknown |
| 63 | IACSP023168 | IAC914216 | Unknown |
| 64 | IACSP042503 | PO8862 | Unknown |
| 65 | IACSP042504 | IACSP955037 | Unknown |
| 66 | IACSP042509 | SP913059 | CT961414 |
| 67 | IACSP042510 | IAC913093 | CT931231 |
| 68 | IACSP043123 | IACSP953028 | RB855453 |
| 69 | IACSP043148 | IACSP953018 | IAC911099 |
| 70 | IACSP043150 | IACSP953018 | CT014455 |
| 71 | IACSP043259 | IACSP966026 | CT961307 |
| 72 | IACSP045081 | IACSP966026 | SP775181 |
| 73 | IACSP046032 | IACSP953028 | CTC957 |

| | | | |
|---|---|---|---|
| 74 | IACSP046035 | SP913059 | SP775181 |
| 75 | IACSP046073 | IACSP953028 | SP701143 |
| 76 | IACSP046077 | IACSP972109 | SP901644 |
| 77 | IACSP046152 | IAC911099 | SP88813 |
| 78 | IACSP933046 | SP791011 | SP803280 |
| 79 | IACSP953018 | SP842189 | CTC9 |
| 80 | IACSP956114 | IAC873187 | CTC9638 |
| 81 | IACSP962008 | SP80144 | SP832847 |
| 82 | IACSP963056 | SP826108 | IACSP933046 |
| 83 | IACSP963069 | SP803280 | IACSP933046 |
| 84 | IACSP963076 | SP847017 | RB855453 |
| 85 | IACSP973384 | RB855113 | IACSP966026 |
| 86 | IACSP974039 | RB835486 | SP924221 |
| 87 | IACSP974048 | SP842066 | IACSP956114 |
| 88 | IACSP982053 | SP847017 | SP801842 |
| 89 | IACSP983011 | IAC913093 | CTC9019 |
| 90 | IACSP993369 | SP901616 | IACCTC059607 |
| 91 | IACSP994011 | SP842025 | RB855453 |
| 92 | IJ76293 | *S. robustum* | IAC863154 |
| 93 | IN8458 | *S. spontaneum* | SP801842 |
| 94 | IN8488 | *S. spontaneum* | Unknown |
| 95 | NG57213 | Unknown | Unknown |
| 96 | RB935744 | RB835089 | TUC717 |
| 97 | SP832847 | HJ5741 | Unknown |

**Table S2. List of genotypes in the validation panels and their origin.** The varieties and elite clones are hybrids that originated from Brazilian sugarcane breeding programs: the Agronomic Institute of Campinas (IAC), the Inter-University Network for the Development of the Sugarcane Sector (RIDESA), and the Sugarcane Research Center (CTC).

| ID Number | Genotype | Panel | Origin |
|---|---|---|---|
| 1 | Ajax | Wild | F2 (*S. officinarum*) |
| 2 | Badilla | Wild | *S. officinarum* x NG96 |
| 3 | Badilla de Java | Wild | F1 (*S. officinarum*) |
| 4 | Caiana | Wild | F1 (*S. officinarum*) |
| 5 | Caiana fita | Wild | F1 (*S. officinarum*) |
| 6 | Caiana riscada | Wild | F1 (*S. officinarum*) |
| 7 | Ceram red | Wild | *S. officinarum* |
| 8 | Chin | Wild | *S. barberi* x ? |
| 9 | Chunnee | Wild | *S. barberi* |
| 10 | CTC9 | Variety/Elite | CTC variety |
| 11 | Formosa 4 | Wild | F1 (*S. officinarum* x ?) |
| 12 | Gandacheni | Wild | *S. barberi* x ? |
| 13 | IAC52150 | Variety/Elite | IAC elite clone |
| 14 | IAC862210 | Variety/Elite | IAC variety |
| 15 | IAC862480 | Variety/Elite | IAC variety |
| 16 | IAC873396 | Variety/Elite | IAC variety |
| 17 | IAC911099 | Variety/Elite | IAC variety |
| 18 | IAC912195 | Variety/Elite | IAC variety |

| | | | |
|---|---|---|---|
| 19 | IAC912218 | Variety/Elite | IAC variety |
| 20 | IACSP022074 | Variety/Elite | IAC elite clone |
| 21 | IACSP932060 | Variety/Elite | IAC variety |
| 22 | IACSP936006 | Variety/Elite | IAC variety |
| 23 | IACSP942094 | Variety/Elite | IAC variety |
| 24 | IACSP942101 | Variety/Elite | IAC variety |
| 25 | IACSP944004 | Variety/Elite | IAC variety |
| 26 | IACSP951218 | Variety/Elite | IAC variety |
| 27 | IACSP952078 | Variety/Elite | IAC elite clone |
| 28 | IACSP955000 | Variety/Elite | IAC variety |
| 29 | IACSP955094 | Variety/Elite | IAC variety |
| 30 | IACSP961005 | Variety/Elite | IAC elite clone |
| 31 | IACSP962019 | Variety/Elite | IAC elite clone |
| 32 | IACSP962042 | Variety/Elite | IAC variety |
| 33 | IACSP962100 | Variety/Elite | IAC elite clone |
| 34 | IACSP963048 | Variety/Elite | IAC elite clone |
| 35 | IACSP963060 | Variety/Elite | IAC variety |
| 36 | IACSP967506 | Variety/Elite | IAC elite clone |
| 37 | IACSP967569 | Variety/Elite | IAC variety |
| 38 | IACSP967586 | Variety/Elite | IAC elite clone |
| 39 | IACSP972000 | Variety/Elite | IAC elite clone |
| 40 | IACSP972020 | Variety/Elite | IAC elite clone |
| 41 | IACSP972023 | Variety/Elite | IAC elite clone |

| | | | |
|---|---|---|---|
| 42 | IACSP972028 | Variety/Elite | IAC elite clone |
| 43 | IACSP972053 | Variety/Elite | IAC elite clone |
| 44 | IACSP972055 | Variety/Elite | IAC elite clone |
| 45 | IACSP972098 | Variety/Elite | IAC elite clone |
| 46 | IACSP973046 | Variety/Elite | IAC variety |
| 47 | IACSP973313 | Variety/Elite | IAC elite clone |
| 48 | IACSP973331 | Variety/Elite | IAC elite clone |
| 49 | IACSP976682 | Variety/Elite | IAC elite clone |
| 50 | IACSP977018 | Variety/Elite | IAC elite clone |
| 51 | IACSP977065 | Variety/Elite | IAC elite clone |
| 52 | IACSP977074 | Variety/Elite | IAC elite clone |
| 53 | IACSP977077 | Variety/Elite | IAC elite clone |
| 54 | IACSP982030 | Variety/Elite | IAC elite clone |
| 55 | IACSP983020 | Variety/Elite | IAC elite clone |
| 56 | IACSP983021 | Variety/Elite | IAC elite clone |
| 57 | IACSP983099 | Variety/Elite | IAC elite clone |
| 58 | IACSP985010 | Variety/Elite | IAC elite clone |
| 59 | IACSP986202 | Variety/Elite | IAC elite clone |
| 60 | IACSP991305 | Variety/Elite | IAC elite clone |
| 61 | IACSP991308 | Variety/Elite | IAC elite clone |
| 62 | IACSP993085 | Variety/Elite | IAC elite clone |
| 63 | IACSP994007 | Variety/Elite | IAC elite clone |
| 64 | IACSP994008 | Variety/Elite | IAC elite clone |

| | | | |
|---|---|---|---|
| 65 | IACSP994010 | Variety/Elite | IAC elite clone |
| 66 | IACSP994013 | Variety/Elite | IAC elite clone |
| 67 | IJ76313 | Wild | F1 (*S. officinarum* x ?) |
| 68 | IJ76317 | Wild | F1 (*S. officinarum* x ?) |
| 69 | IJ76325 | Wild | F1 (*S. officinarum* x ?) |
| 70 | IJ76560 | Wild | F1 (*S. officinarum* x ?) |
| 71 | IJ76566 | Wild | F1 (*S. officinarum* x ?) |
| 72 | IM76229 | Wild | *S. robustum* |
| 73 | IN84105 | Wild | F1 (*S. officinarum* x ?) |
| 74 | IN84126 | Wild | F1 (*S. officinarum*) |
| 75 | MZ151 | Wild | F1 (*S. officinarum*) |
| 76 | NG2121 | Wild | F1 (*S. officinarum*) |
| 77 | NG5750 | Wild | F1 (*S. officinarum*) |
| 78 | NG7718 | Wild | F1 (*S. officinarum*) |
| 79 | Pitu | Wild | F1 (*S. officinarum*) |
| 80 | RB72454 | Variety/Elite | RIDESA variety |
| 81 | RB835486 | Variety/Elite | RIDESA variety |
| 82 | RB855156 | Variety/Elite | RIDESA variety |
| 83 | RB867515 | Variety/Elite | RIDESA variety |
| 84 | RB925211 | Variety/Elite | RIDESA variety |
| 85 | Sabura | Wild | Hybrid (*S. officinarum* x ?) |
| 86 | SES205A | Wild | *S. spontaneum* |
| 87 | SP701143 | Variety/Elite | CTC variety |

| | | | |
|---|---|---|---|
| 88 | SP791011 | Variety/Elite | CTC variety |
| 89 | SP803280 | Variety/Elite | CTC variety |
| 90 | SP8642 | Variety/Elite | CTC variety |
| 91 | US1008 | Wild | *S. spontaneum* x US60313 |
| 92 | White transparent | Wild | *S. officinarum* |

**Table S3. Markers significantly associated with sugarcane mosaic virus (SCMV) resistance by genome-wide association mapping.** For each marker–trait association, the chromosome location, position, marker-effect model used, -$\log_{10}$(p) value, effect, $r^2$ value and polymorphism (reference/alternative alleles) are provided. $r^2$ values and effects are not provided for those of the diploidized general model.

| Marker | Chromosome | Position | Model | -$\log_{10}$(p) | Effect | $r^2$ | Polymorphism |
|---|---|---|---|---|---|---|---|
| Chr1A_111976277 | 1 | 111976277 | 1-dom-ref | 5.67 | 10.6 | 0.267 | C/T |
| Chr1A_111976277 | 1 | 111976277 | Diplo-additive | 5.67 | 10.6 | 0.267 | C/T |
| Chr1A_111976277 | 1 | 111976277 | Diplo-general | 5.67 | NA | 0.267 | C/T |
| Chr3A_44961867 | 3 | 44961867 | 1-dom-ref | 7.75 | 15.9 | 0.062 | C/T |
| Chr3A_44961867 | 3 | 44961867 | Diplo-general | 6.94 | NA | 0.062 | C/T |
| Chr3A_5054269 | 3 | 5054269 | Diplo-general | 5.89 | NA | 0.071 | A/G |
| Chr4A_64827729 | 4 | 64827729 | 1-dom-ref | 5.56 | 12.8 | 0.066 | A/G |
| Chr4A_64827729 | 4 | 64827729 | Diplo-additive | 5.56 | 12.8 | 0.066 | A/G |
| Chr4A_64827729 | 4 | 64827729 | Diplo-general | 5.56 | NA | 0.066 | A/G |
| Chr5A_2059258 | 5 | 2059258 | 1-dom-ref | 11.08 | 12.7 | 0.158 | A/T |

| | | | | | | | |
|---|---|---|---|---|---|---|---|
| Chr5A_2059258 | 5 | 2059258 | Diplo-additive | 11.08 | 12.7 | 0.158 | A/T |
| Chr5A_2059258 | 5 | 2059258 | Diplo-general | 11.08 | NA | 0.158 | A/T |
| Chr5A_2360726 | 5 | 2360726 | 1-dom-ref | 9.96 | 18.7 | 0.145 | C/A |
| Chr5A_2360726 | 5 | 2360726 | Diplo-additive | 9.96 | 18.7 | 0.145 | C/A |
| Chr5A_2360726 | 5 | 2360726 | Diplo-general | 9.96 | NA | 0.145 | C/A |
| Chr5A_79992188 | 5 | 79992188 | Diplo-general | 6.36 | NA | 0.170 | G/A |
| Chr7A_33360091 | 7 | 33360091 | 1-dom-ref | 5.78 | 9.33 | 0.299 | G/A |
| Chr7A_33360091 | 7 | 33360091 | Diplo-additive | 5.78 | 9.33 | 0.299 | G/A |
| Chr7A_33360091 | 7 | 33360091 | Diplo-general | 5.78 | NA | 0.299 | G/A |
| Contigs_3441998 | - | 3441998 | Diplo-general | 6.36 | NA | 0.017 | C/T |

**Table S4. $r^2$ values of significant marker–trait associations identified by by genome-wide association mapping in the validation panels.** The values are provided for different models (diploidized (diplo) and simplex dominant reference (1-dom-ref)) and are separate for the wild accessions and the variety and elite clone panels.

| Marker | Model | $r^2$ | |
|---|---|---|---|
| | | Wild | Variety/Elite |
| Chr1A_111976277 | Diplo | 0.004 | 0.010 |
| Chr1A_111976277 | 1-dom-ref | 0.004 | 0.034 |
| Chr3A_5054269 | Diplo | 0.016 | 0.236 |
| Chr3A_5054269 | 1-dom-ref | 0.000 | 0.000 |
| Chr4A_64827729 | Diplo | 0.004 | 0.000 |
| Chr4A_64827729 | 1-dom-ref | 0.000 | 0.000 |
| Chr5A_2360726 | Diplo | 0.013 | 0.020 |
| Chr5A_2360726 | 1-dom-ref | 0.000 | 0.000 |
| Chr5A_79992188 | Diplo | 0.014 | 0.046 |
| Chr5A_79992188 | 1-dom-ref | 0.000 | 0.000 |
| Chr7A_33360091 | Diplo | 0.010 | 0.149 |
| Chr7A_33360091 | 1-dom-ref | 0.000 | 0.000 |
| Contigs_3441998 | Diplo | 0.002 | 0.005 |
| Contigs_3441998 | 1-dom-ref | 0.000 | 0.000 |

**Table S5. Number and percentage of conserved orthologs from Viridiplantae found in the complete transcriptome assembly.**

| BUSCOs | Number (%) |
|---|---|
| **Complete and single copy** | 110 (25.9%) |
| **Complete and duplicated** | 246 (57.9%) |
| **Fragmented** | 63 (14.8%) |
| **Missing** | 6 (1.4%) |
| **Total searched** | 425 |

**Table S6. Annotations of markers associated with sugarcane mosaic virus (SCMV) resistance.** For each marker, the transcript, network module to which the corresponding gene belongs, alignment percentage of identity (% ID), alignment E-value, distance between the marker and gene, and Gene Ontology annotation information are provided.

| Marker | Transcript | Module | % ID | E-Value | Distance | Annotation |
|---|---|---|---|---|---|---|
| Chr1A_101288487 | TRINITY_DN100313_c0_g1_i1 | NA | 96.906 | 0 | 1177 | - |
| Chr1A_11663583 | TRINITY_DN22017_c0_g1_i4 | 2 | 96.667 | 1.40E-79 | 300 | - |
| Chr1A_11663583 | TRINITY_DN22017_c0_g1_i5 | 2 | 94.828 | 1.29E-179 | 632 | - |
| Chr1A_11663583 | TRINITY_DN22017_c0_g1_i6 | 2 | 96.667 | 1.27E-79 | 300 | - |
| Chr1A_11663583 | TRINITY_DN22017_c0_g1_i7 | 2 | 95.198 | 0 | 758 | - |
| Chr1A_11663583 | TRINITY_DN22017_c0_g1_i1 | 2 | 94.236 | 2.14E-172 | 608 | - |
| Chr1A_11663583 | TRINITY_DN22017_c0_g1_i2 | 2 | 95.198 | 0 | 756 | - |
| Chr1A_11663583 | TRINITY_DN22017_c0_g1_i3 | 2 | 94.776 | 7.43E-52 | 207 | - |
| Chr1A_11663583 | TRINITY_DN22017_c0_g1_i8 | 2 | 94.776 | 7.43E-52 | 207 | - |
| Chr1A_11663583 | TRINITY_DN22017_c0_g1_i9 | 2 | 93.734 | 4.64E-169 | 597 | - |
| Chr1A_14820959 | TRINITY_DN105877_c0_g1_i1 | 1 | 96.797 | 7.84E-131 | 470 | - |
| Chr1A_14820959 | TRINITY_DN130_c0_g1_i4 | 61 | 93.508 | 0 | 1053 | - |
| Chr1A_14820959 | TRINITY_DN130_c0_g1_i7 | 61 | 94.643 | 0 | 680 | - |
| Chr1A_14820959 | TRINITY_DN130_c0_g1_i2 | 61 | 94.643 | 0 | 680 | - |
| Chr1A_14820959 | TRINITY_DN130_c0_g1_i9 | 61 | 93.508 | 0 | 1053 | - |
| Chr1A_14820959 | TRINITY_DN130_c0_g1_i23 | 61 | 93.508 | 0 | 1053 | - |
| Chr1A_14820959 | TRINITY_DN130_c0_g1_i20 | 61 | 94.643 | 0 | 680 | - |

| | | | | | | |
|---|---|---|---|---|---|---|
| Chr1A_14820959 | TRINITY_DN130_c0_g1_i16 | 61 | 94.643 | 0 | 680 | - |
| Chr1A_14820959 | TRINITY_DN20299_c0_g3_i1 | NA | 95.827 | 0 | 1044 | - |
| Chr1A_21076224 | TRINITY_DN22610_c0_g1_i1 | 1 | 99.209 | 7.68E-127 | 457 | - |
| Chr1A_21076224 | TRINITY_DN22610_c0_g1_i2 | 1 | 95.885 | 3.10E-107 | 392 | - |
| Chr1A_21076224 | TRINITY_DN22610_c0_g1_i3 | 1 | 96.838 | 3.98E-116 | 422 | - |
| Chr1A_21076224 | TRINITY_DN6805_c0_g2_i7 | 54 | 97.837 | 0 | 719 | UPF0301 protein CT0663 |
| Chr1A_21076224 | TRINITY_DN6805_c0_g2_i6 | 54 | 98.798 | 0 | 741 | UPF0301 protein CT0663 |
| Chr1A_32004665 | TRINITY_DN81589_c0_g1_i4 | NA | 93.499 | 0 | 1526 | - |
| Chr1A_32004665 | TRINITY_DN81589_c0_g1_i6 | NA | 93.321 | 0 | 1517 | - |
| Chr1A_32004665 | TRINITY_DN81589_c0_g1_i7 | NA | 93.321 | 0 | 1517 | - |
| Chr1A_32004665 | TRINITY_DN81589_c0_g1_i1 | NA | 95 | 3.42E-111 | 407 | - |
| Chr1A_32004665 | TRINITY_DN81589_c0_g1_i2 | NA | 93.499 | 0 | 1526 | - |
| Chr1A_32004665 | TRINITY_DN81589_c0_g1_i3 | NA | 95 | 2.17E-111 | 407 | - |
| Chr1A_32004665 | TRINITY_DN81589_c0_g1_i8 | NA | 95 | 3.45E-111 | 407 | - |
| Chr1A_32004665 | TRINITY_DN81589_c0_g1_i9 | NA | 95 | 2.15E-111 | 407 | - |
| Chr1A_32004665 | TRINITY_DN81589_c0_g1_i12 | NA | 92.243 | 0 | 817 | - |
| Chr1A_33266946 | TRINITY_DN12969_c0_g2_i3 | 2 | 97.924 | 4.95E-140 | 155 | - |
| Chr1A_33266946 | TRINITY_DN123250_c0_g1_i4 | 9 | 93.75 | 8.65E-89 | 1065 | - |
| Chr1A_64093150 | TRINITY_DN94565_c0_g1_i5 | NA | 98.987 | 0 | 706 | - |
| Chr1A_64093150 | TRINITY_DN94565_c0_g1_i6 | NA | 99.031 | 0 | 739 | - |
| Chr1A_74790184 | TRINITY_DN3672_c0_g1_i24 | 10 | 92.555 | 0 | 961 | Alpha-galactosidase 1 |
| Chr1A_74790184 | TRINITY_DN3672_c0_g1_i8 | 10 | 92.555 | 0 | 961 | Alpha-galactosidase 1 |

| | | | | | | |
|---|---|---|---|---|---|---|
| Chr1A_74790184 | TRINITY_DN3672_c0_g1_i23 | 10 | 96.171 | 0 | 1099 | Alpha-galactosidase |
| Chr1A_74790184 | TRINITY_DN3672_c0_g1_i22 | 10 | 90.011 | 0 | 1103 | Alpha-galactosidase 1 |
| Chr1A_74790184 | TRINITY_DN3672_c0_g1_i26 | 10 | 96.495 | 0 | 795 | Alpha-galactosidase |
| Chr1A_74790184 | TRINITY_DN3672_c0_g1_i16 | 10 | 90.011 | 0 | 1103 | Alpha-galactosidase 1 |
| Chr1A_74790184 | TRINITY_DN3672_c0_g1_i14 | 10 | 90.011 | 0 | 1103 | Alpha-galactosidase 1 |
| Chr1A_74790184 | TRINITY_DN3672_c0_g1_i15 | 10 | 90.011 | 0 | 1103 | Alpha-galactosidase 1 |
| Chr1A_74790184 | TRINITY_DN3672_c0_g1_i13 | 10 | 90.011 | 0 | 1103 | Alpha-galactosidase |
| Chr1A_90316612 | TRINITY_DN73_c2_g1_i3 | 20 | 94.629 | 0 | 1554 | Disease resistance protein RGA5 |
| Chr1A_90316612 | TRINITY_DN73_c2_g1_i1 | 20 | 99.119 | 0 | 1834 | Disease resistance protein RGA5 |
| Chr1A_90316612 | TRINITY_DN73_c2_g1_i6 | 20 | 94.629 | 0 | 1554 | Nuclear transcription factor Y subunit A-7 |
| Chr1A_90316612 | TRINITY_DN73_c2_g1_i4 | 20 | 99.119 | 0 | 1834 | Nuclear transcription factor Y subunit A-7 |
| Chr2A_103190628 | TRINITY_DN58652_c0_g1_i4 | NA | 98.926 | 0 | 1164 | - |
| Chr2A_103190628 | TRINITY_DN58652_c0_g1_i1 | NA | 99.455 | 0 | 665 | - |
| Chr2A_103190628 | TRINITY_DN58652_c0_g1_i2 | NA | 98.76 | 0 | 861 | - |
| Chr2A_103190628 | TRINITY_DN58652_c0_g1_i3 | NA | 98.971 | 0 | 1044 | - |
| Chr2A_1809005 | TRINITY_DN1924_c0_g2_i1 | NA | 95.448 | 0 | 1173 | - |
| Chr2A_22381106 | TRINITY_DN102055_c1_g2_i1 | NA | 92.368 | 1.25E-149 | 1277 | - |
| Chr2A_22381106 | TRINITY_DN67121_c0_g1_i1 | NA | 91.165 | 0 | 1799 | - |
| Chr2A_23071827 | TRINITY_DN2434_c0_g1_i7 | 57 | 99.107 | 1.47E-109 | 403 | Tetratricopeptide repeat domain-containing protein PYG7, chloroplastic |
| Chr2A_23071827 | TRINITY_DN2434_c0_g1_i8 | 57 | 96.558 | 0 | 1709 | Tetratricopeptide repeat domain-containing protein PYG7, chloroplastic |
| Chr2A_23071827 | TRINITY_DN2434_c0_g1_i16 | 57 | 97.975 | 0 | 1363 | Tetratricopeptide repeat domain-containing protein PYG7, chloroplastic |

| | | | | | | |
|---|---|---|---|---|---|---|
| Chr2A_23071827 | TRINITY_DN2434_c0_g1_i13 | 57 | 96.558 | 0 | 1709 | Tetratricopeptide repeat domain-containing protein PYG7, chloroplastic |
| Chr2A_24874343 | TRINITY_DN25150_c0_g3_i1 | NA | 93.455 | 1.25E-159 | 1476 | - |
| Chr2A_59276886 | TRINITY_DN10411_c0_g1_i5 | NA | 97.844 | 0 | 1275 | Protein upstream of FLC |
| Chr2A_59276886 | TRINITY_DN10411_c0_g1_i6 | NA | 99.367 | 2.67E-74 | 287 | Protein upstream of FLC |
| Chr2A_59276886 | TRINITY_DN10411_c0_g1_i2 | NA | 99.367 | 2.63E-74 | 287 | Protein upstream of FLC |
| Chr2A_59276886 | TRINITY_DN10411_c0_g1_i8 | NA | 97.844 | 0 | 1275 | Protein upstream of FLC |
| Chr2A_59276886 | TRINITY_DN49441_c0_g1_i4 | NA | 96.688 | 0 | 1783 | Protein upstream of FLC |
| Chr2A_59276886 | TRINITY_DN49441_c0_g1_i6 | NA | 96.688 | 0 | 1783 | Protein upstream of FLC |
| Chr2A_59276886 | TRINITY_DN49441_c0_g1_i1 | NA | 96.609 | 0 | 1687 | Protein upstream of FLC |
| Chr2A_59276886 | TRINITY_DN49441_c0_g1_i2 | NA | 96.688 | 0 | 1783 | Protein upstream of FLC |
| Chr2A_59276886 | TRINITY_DN49441_c0_g1_i8 | NA | 99.154 | 0 | 848 | Protein upstream of FLC |
| Chr2A_59276886 | TRINITY_DN49441_c0_g1_i9 | NA | 99.154 | 0 | 848 | Protein upstream of FLC |
| Chr2A_59276886 | TRINITY_DN49441_c0_g1_i16 | NA | 99.154 | 0 | 848 | Protein upstream of FLC |
| Chr2A_59276886 | TRINITY_DN49441_c0_g1_i15 | NA | 96.702 | 0 | 1692 | Protein upstream of FLC |
| Chr2A_59276886 | TRINITY_DN49441_c0_g1_i11 | NA | 96.609 | 0 | 1687 | Protein upstream of FLC |
| Chr2A_77406645 | TRINITY_DN36993_c0_g3_i3 | 23 | 97.304 | 0 | 534 | - |
| Chr2A_85950136 | TRINITY_DN2040_c0_g1_i4 | 14 | 98.814 | 2.34E-124 | 451 | - |
| Chr2A_85950136 | TRINITY_DN2040_c0_g1_i6 | 14 | 98.814 | 1.33E-124 | 451 | - |
| Chr2A_85950136 | TRINITY_DN2040_c0_g1_i3 | 14 | 98.814 | 2.35E-124 | 451 | - |
| Chr2A_85950136 | TRINITY_DN2040_c0_g1_i8 | 14 | 98.814 | 1.22E-124 | 451 | - |
| Chr2A_85950136 | TRINITY_DN2040_c0_g1_i10 | 14 | 98.814 | 2.35E-124 | 451 | - |
| Chr2A_94947162 | TRINITY_DN176071_c0_g1_i5 | NA | 96.864 | 7.46E-134 | 481 | - |

| | | | | | | |
|---|---|---|---|---|---|---|
| Chr2A_94947162 | TRINITY_DN176071_c0_g1_i1 | NA | 96.864 | 5.45E-134 | 481 | - |
| Chr3A_14502333 | TRINITY_DN133571_c0_g1_i1 | 3 | 97.849 | 2.44E-86 | 322 | - |
| Chr3A_15810035 | TRINITY_DN18724_c0_g2_i1 | NA | 99.065 | 0 | 1533 | - |
| Chr3A_15810035 | TRINITY_DN18724_c0_g2_i3 | NA | 94.488 | 1.43E-161 | 573 | - |
| Chr3A_15810035 | TRINITY_DN18724_c0_g2_i2 | NA | 96.653 | 0 | 1227 | - |
| Chr3A_15810035 | TRINITY_DN18724_c0_g2_i5 | NA | 97.471 | 0 | 1469 | - |
| Chr3A_15810035 | TRINITY_DN18724_c0_g2_i4 | NA | 98.499 | 0 | 1291 | - |
| Chr3A_15810035 | TRINITY_DN18724_c0_g2_i7 | NA | 96.429 | 0 | 817 | - |
| Chr3A_15810035 | TRINITY_DN18724_c0_g2_i6 | NA | 98.093 | 0 | 638 | - |
| Chr3A_15810035 | TRINITY_DN52807_c3_g1_i2 | NA | 97.108 | 0 | 1382 | - |
| Chr3A_15810035 | TRINITY_DN52807_c3_g1_i1 | NA | 98.897 | 0 | 1452 | - |
| Chr3A_18503343 | TRINITY_DN81920_c0_g1_i2 | 7 | 94.195 | 0 | 300 | - |
| Chr3A_18503343 | TRINITY_DN1246_c1_g1_i9 | 47 | 99.484 | 0 | 463 | Serine/threonine-protein phosphatase PP2A-4 catalytic subunit |
| Chr3A_19597062 | TRINITY_DN552_c1_g1_i5 | 49 | 99.277 | 0 | 1249 | Chaperone protein dnaJ C76, chloroplastic |
| Chr3A_19597062 | TRINITY_DN552_c1_g1_i7 | 49 | 99.277 | 0 | 1249 | Chaperone protein dnaJ C76, chloroplastic |
| Chr3A_19597062 | TRINITY_DN552_c1_g1_i15 | 49 | 99.439 | 0 | 1618 | Chaperone protein dnaJ C76, chloroplastic |
| Chr3A_19597062 | TRINITY_DN552_c1_g1_i14 | 49 | 99.439 | 0 | 1618 | Chaperone protein dnaJ C76, chloroplastic |
| Chr3A_19597062 | TRINITY_DN552_c1_g1_i11 | 49 | 99.277 | 0 | 1249 | Chaperone protein dnaJ C76, chloroplastic |
| Chr3A_19597062 | TRINITY_DN552_c1_g1_i10 | 49 | 99.277 | 0 | 1249 | Chaperone protein dnaJ C76, chloroplastic |
| Chr3A_27409051 | TRINITY_DN32141_c2_g1_i1 | 2 | 95.5 | 3.19E-85 | 88 | - |
| Chr3A_33621969 | TRINITY_DN27357_c0_g1_i2 | 3 | 90.291 | 3.77E-70 | 694 | - |
| Chr3A_33621969 | TRINITY_DN24045_c0_g2_i18 | NA | 98.525 | 0 | 994 | Protein pigment defective 338, chloroplastic |

| | | | | | | |
|---|---|---|---|---|---|---|
| Chr3A_5040761 | TRINITY_DN36461_c0_g1_i1 | 1 | 95.858 | 5.46E-154 | 293 | - |
| Chr3A_5040761 | TRINITY_DN172053_c0_g1_i1 | NA | 96.507 | 1.36E-103 | 1074 | - |
| Chr3A_5040795 | TRINITY_DN36461_c0_g1_i1 | 1 | 95.858 | 5.46E-154 | 327 | - |
| Chr3A_5040795 | TRINITY_DN172053_c0_g1_i1 | NA | 96.507 | 1.36E-103 | 1040 | - |
| Chr3A_5054269 | TRINITY_DN901_c0_g1_i5 | 6 | 90.251 | 8.57E-127 | 460 | Pentatricopeptide repeat-containing protein At3g22670, mitochondrial |
| Chr3A_5054269 | TRINITY_DN901_c0_g1_i6 | 6 | 90.251 | 5.12E-127 | 460 | Pentatricopeptide repeat-containing protein At3g22670, mitochondrial |
| Chr3A_5054269 | TRINITY_DN901_c0_g1_i20 | 6 | 90.251 | 7.45E-127 | 460 | Pentatricopeptide repeat-containing protein At3g22670, mitochondrial |
| Chr3A_5054269 | TRINITY_DN901_c0_g1_i16 | 6 | 90.251 | 4.95E-127 | 460 | Pentatricopeptide repeat-containing protein At3g22670, mitochondrial |
| Chr3A_5054269 | TRINITY_DN901_c0_g1_i17 | 6 | 90.251 | 9.44E-127 | 460 | Pentatricopeptide repeat-containing protein At3g22670, mitochondrial |
| Chr3A_5054269 | TRINITY_DN901_c0_g1_i15 | 6 | 90.251 | 9.54E-127 | 460 | Pentatricopeptide repeat-containing protein At3g22670, mitochondrial |
| Chr3A_5054269 | TRINITY_DN901_c0_g1_i12 | 6 | 90.251 | 7.28E-127 | 460 | Pentatricopeptide repeat-containing protein At3g22670, mitochondrial |
| Chr3A_5054269 | TRINITY_DN901_c0_g1_i13 | 6 | 90.251 | 4.43E-127 | 460 | Pentatricopeptide repeat-containing protein At3g22670, mitochondrial |
| Chr3A_57785310 | TRINITY_DN1485_c2_g1_i2 | NA | 97.18 | 0 | 894 | - |
| Chr3A_57785310 | TRINITY_DN1485_c2_g1_i3 | NA | 97.556 | 0 | 905 | - |
| Chr3A_57785310 | TRINITY_DN1485_c2_g1_i1 | NA | 97.804 | 0 | 859 | - |
| Chr3A_59179238 | TRINITY_DN3857_c1_g1_i9 | 20 | 99.266 | 0 | 941 | SWI/SNF complex subunit SWI3C |
| Chr3A_6428925 | TRINITY_DN104159_c0_g3_i1 | NA | 98.165 | 0 | 1092 | - |
| Chr3A_67612941 | TRINITY_DN22619_c1_g2_i3 | 1 | 98.84 | 0 | 547 | - |

| | | | | | | |
|---|---|---|---|---|---|---|
| Chr4A_11325067 | TRINITY_DN23211_c0_g1_i4 | 9 | 95.789 | 8.51E-82 | 1086 | - |
| Chr4A_11325067 | TRINITY_DN6374_c0_g1_i7 | 13 | 95 | 0 | 1428 | General transcription factor 3C polypeptide 2 |
| Chr4A_15495314 | TRINITY_DN4216_c0_g1_i4 | 41 | 97.887 | 1.51E-62 | 246 | Transcription factor bHLH63 |
| Chr4A_15495314 | TRINITY_DN4216_c0_g1_i18 | 41 | 97.887 | 1.55E-62 | 246 | Transcription factor bHLH63 |
| Chr4A_15495314 | TRINITY_DN4216_c0_g1_i14 | 41 | 97.887 | 1.43E-62 | 246 | Carbonic anhydrase, chloroplastic |
| Chr4A_15495314 | TRINITY_DN4216_c0_g1_i11 | 41 | 97.887 | 1.47E-62 | 246 | Carbonic anhydrase, chloroplastic |
| Chr4A_62003722 | TRINITY_DN4047_c0_g1_i3 | 22 | 93.671 | 3.44E-60 | 785 | - |
| Chr4A_62003722 | TRINITY_DN1484_c0_g1_i5 | 24 | 94.451 | 0 | 1167 | Very-long-chain aldehyde decarbonylase GL1-2 |
| Chr4A_67754808 | TRINITY_DN134539_c0_g1_i1 | NA | 95.249 | 0 | 23 | - |
| Chr4A_67754808 | TRINITY_DN156943_c0_g1_i1 | NA | 92 | 4.62E-130 | 1876 | - |
| Chr5A_14593032 | TRINITY_DN131549_c0_g1_i3 | NA | 97.044 | 3.55E-91 | 1041 | SUMO-conjugating enzyme SCE1 |
| Chr5A_14593032 | TRINITY_DN131549_c0_g1_i2 | NA | 99.283 | 2.01E-140 | 702 | SUMO-conjugating enzyme SCE1 |
| Chr5A_14660425 | TRINITY_DN116305_c0_g1_i1 | NA | 92.123 | 1.30E-110 | 438 | - |
| Chr5A_14660425 | TRINITY_DN144282_c0_g1_i1 | NA | 95.122 | 0 | 1760 | - |
| Chr5A_14660427 | TRINITY_DN116305_c0_g1_i1 | NA | 92.123 | 1.30E-110 | 436 | - |
| Chr5A_14660427 | TRINITY_DN144282_c0_g1_i1 | NA | 95.122 | 0 | 1762 | - |
| Chr5A_16353126 | TRINITY_DN4434_c0_g1_i6 | 47 | 96.838 | 0 | 852 | - |
| Chr5A_2059258 | TRINITY_DN93396_c0_g1_i4 | NA | 91.777 | 0 | 125 | - |
| Chr5A_2360726 | TRINITY_DN1781_c0_g1_i5 | 43 | 97.664 | 0 | 1458 | Probable allantoinase |
| Chr5A_2360726 | TRINITY_DN1781_c0_g1_i7 | 43 | 99.213 | 2.47E-57 | 228 | Probable allantoinase |
| Chr5A_2360726 | TRINITY_DN1781_c0_g1_i1 | 43 | 99.213 | 9.71E-57 | 228 | Probable allantoinase |
| Chr5A_2360726 | TRINITY_DN1781_c0_g1_i2 | 43 | 99.078 | 4.17E-105 | 388 | Probable allantoinase |

| | | | | | | |
|---|---|---|---|---|---|---|
| Chr5A_2360726 | TRINITY_DN1781_c0_g1_i3 | 43 | 99.213 | 6.32E-57 | 228 | Probable allantoinase |
| Chr5A_2360726 | TRINITY_DN1781_c0_g1_i8 | 43 | 99.213 | 9.45E-57 | 228 | Probable allantoinase |
| Chr5A_2360726 | TRINITY_DN1781_c0_g1_i9 | 43 | 97.664 | 0 | 1458 | Probable allantoinase |
| Chr5A_2360726 | TRINITY_DN1781_c0_g1_i18 | 43 | 99.078 | 2.88E-105 | 388 | Probable allantoinase |
| Chr5A_2360726 | TRINITY_DN1781_c0_g1_i19 | 43 | 97.664 | 0 | 1458 | Probable allantoinase |
| Chr5A_2360726 | TRINITY_DN1781_c0_g1_i16 | 43 | 99.213 | 4.66E-57 | 228 | Probable allantoinase |
| Chr5A_2360726 | TRINITY_DN1781_c0_g1_i15 | 43 | 99.213 | 2.49E-57 | 228 | Probable allantoinase |
| Chr5A_2360726 | TRINITY_DN1781_c0_g1_i12 | 43 | 99.213 | 9.27E-57 | 228 | Probable allantoinase |
| Chr5A_2360726 | TRINITY_DN1781_c0_g1_i13 | 43 | 99.078 | 4.36E-105 | 388 | Probable allantoinase |
| Chr5A_2360726 | TRINITY_DN1781_c0_g1_i10 | 43 | 99.078 | 4.28E-105 | 388 | Probable allantoinase |
| Chr5A_2360726 | TRINITY_DN1781_c0_g1_i11 | 43 | 99.213 | 9.48E-57 | 228 | Probable allantoinase |
| Chr5A_30390140 | TRINITY_DN263_c0_g1_i4 | 1 | 95.749 | 0 | 867 | - |
| Chr5A_30390140 | TRINITY_DN263_c0_g1_i5 | 1 | 95.387 | 0 | 856 | - |
| Chr5A_30390140 | TRINITY_DN263_c0_g1_i7 | 1 | 95.387 | 0 | 856 | - |
| Chr5A_30390140 | TRINITY_DN263_c0_g1_i8 | 1 | 95.135 | 0 | 867 | - |
| Chr5A_30390140 | TRINITY_DN263_c0_g1_i9 | 1 | 95.387 | 0 | 856 | - |
| Chr5A_30390140 | TRINITY_DN263_c0_g1_i16 | 1 | 95.113 | 0 | 894 | - |
| Chr5A_30390140 | TRINITY_DN263_c0_g1_i17 | 1 | 95.387 | 0 | 856 | - |
| Chr5A_30390140 | TRINITY_DN263_c0_g1_i14 | 1 | 95.113 | 0 | 894 | - |
| Chr5A_30390140 | TRINITY_DN263_c0_g1_i15 | 1 | 95.387 | 0 | 856 | - |
| Chr5A_33653357 | TRINITY_DN13062_c0_g1_i6 | 5 | 99.128 | 0 | 1335 | Uncharacterized WD repeat-containing protein C3H5.08c |
| Chr5A_72669419 | TRINITY_DN9643_c0_g1_i8 | 5 | 97.176 | 0 | 1308 | - |

| | | | | | | |
|---|---|---|---|---|---|---|
| Chr5A_77270454 | TRINITY_DN108042_c0_g1_i1 | 1 | 95.902 | 1.59E-108 | 332 | - |
| Chr5A_77270454 | TRINITY_DN3896_c0_g1_i1 | 43 | 92.818 | 4.93E-144 | 91 | - |
| Chr5A_79992188 | TRINITY_DN1397_c0_g1_i4 | 35 | 94.386 | 0 | 1423 | 60S ribosomal protein L3 |
| Chr5A_79992188 | TRINITY_DN1397_c0_g1_i5 | 35 | 97.888 | 0 | 1633 | 60S ribosomal protein L3 |
| Chr5A_79992188 | TRINITY_DN1397_c0_g1_i6 | 35 | 94.174 | 0 | 1410 | 60S ribosomal protein L3 |
| Chr5A_79992188 | TRINITY_DN1397_c0_g1_i7 | 35 | 97.888 | 0 | 1633 | 60S ribosomal protein L3 |
| Chr5A_79992188 | TRINITY_DN1397_c0_g1_i1 | 35 | 98.793 | 0 | 881 | 60S ribosomal protein L3 |
| Chr5A_79992188 | TRINITY_DN1397_c0_g1_i2 | 35 | 94.386 | 0 | 1423 | 60S ribosomal protein L3 |
| Chr5A_79992188 | TRINITY_DN1397_c0_g1_i3 | 35 | 97.888 | 0 | 1633 | 60S ribosomal protein L3 |
| Chr6A_101032231 | TRINITY_DN30376_c0_g1_i4 | 5 | 94.961 | 3.20E-109 | 401 | Putative ribonuclease H protein At1g65750 |
| Chr6A_101032231 | TRINITY_DN30376_c0_g1_i3 | 5 | 94.961 | 2.05E-109 | 401 | - |
| Chr6A_101032276 | TRINITY_DN30376_c0_g1_i4 | 5 | 94.961 | 3.20E-109 | 401 | Putative ribonuclease H protein At1g65750 |
| Chr6A_101032276 | TRINITY_DN30376_c0_g1_i3 | 5 | 94.961 | 2.05E-109 | 401 | - |
| Chr6A_101033559 | TRINITY_DN54972_c0_g1_i4 | NA | 97.328 | 0 | 1197 | - |
| Chr6A_101033559 | TRINITY_DN54972_c0_g1_i6 | NA | 97.328 | 0 | 1197 | - |
| Chr6A_101033559 | TRINITY_DN54972_c0_g1_i1 | NA | 94.432 | 0 | 1986 | - |
| Chr6A_101033559 | TRINITY_DN54972_c1_g1_i1 | NA | 99.659 | 0 | 1609 | - |
| Chr6A_22188379 | TRINITY_DN114411_c0_g1_i3 | NA | 92.78 | 0 | 69 | - |
| Chr6A_22188379 | TRINITY_DN72562_c0_g1_i1 | NA | 90.678 | 4.20E-129 | 1993 | - |
| Chr6A_24185578 | TRINITY_DN73382_c0_g1_i3 | 1 | 98.876 | 0 | 636 | Heavy metal-associated isoprenylated plant protein 21 |
| Chr6A_86163774 | TRINITY_DN69187_c0_g1_i4 | NA | 97.11 | 2.18E-77 | 940 | Peroxisomal (S)-2-hydroxy-acid oxidase GLO5 |
| Chr7A_13200334 | TRINITY_DN22336_c0_g1_i4 | 1 | 95.588 | 3.18E-87 | 1024 | - |

| | | | | | | |
|---|---|---|---|---|---|---|
| Chr7A_26774786 | TRINITY_DN7654_c0_g1_i19 | 1 | 93.725 | 1.56E-103 | 383 | Beta-glucosidase 20 |
| Chr7A_26774786 | TRINITY_DN7654_c0_g1_i15 | 1 | 93.725 | 1.47E-103 | 383 | Beta-glucosidase 20 |
| Chr7A_26774786 | TRINITY_DN7654_c0_g1_i1 | 1 | 92.473 | 0 | 1303 | Beta-glucosidase 20 |
| Chr7A_26774786 | TRINITY_DN7654_c0_g1_i2 | 1 | 93.322 | 0 | 1653 | Beta-glucosidase 20 |
| Chr7A_26774786 | TRINITY_DN7654_c0_g1_i8 | 1 | 91.914 | 0 | 1149 | Beta-glucosidase 20 |
| Chr7A_26774786 | TRINITY_DN7654_c0_g1_i21 | 1 | 92.473 | 0 | 1303 | Beta-glucosidase 20 |
| Chr7A_26774786 | TRINITY_DN7654_c0_g1_i20 | 1 | 91.914 | 0 | 1149 | Beta-glucosidase 20 |
| Chr7A_26774786 | TRINITY_DN7654_c0_g1_i26 | 1 | 93.725 | 1.59E-103 | 383 | Beta-glucosidase 20 |
| Chr7A_26774786 | TRINITY_DN7654_c0_g1_i25 | 1 | 97.561 | 1.03E-72 | 279 | Beta-glucosidase 20 |
| Chr7A_26774786 | TRINITY_DN7654_c0_g1_i24 | 1 | 91.914 | 0 | 1149 | Beta-glucosidase 20 |
| Chr7A_26774786 | TRINITY_DN7654_c0_g1_i28 | 1 | 93.725 | 8.87E-104 | 383 | Beta-glucosidase 20 |
| Chr7A_33360091 | TRINITY_DN131579_c0_g1_i1 | NA | 94.639 | 0 | 664 | - |
| Chr7A_33360091 | TRINITY_DN131579_c0_g1_i2 | NA | 94.05 | 0 | 662 | - |
| Chr7A_38817863 | TRINITY_DN82684_c0_g1_i4 | 1 | 95.833 | 6.26E-118 | 539 | - |
| Chr7A_38817863 | TRINITY_DN16706_c0_g2_i1 | NA | 96.396 | 2.33E-98 | 1854 | - |
| Chr7A_65125262 | TRINITY_DN125649_c1_g1_i1 | NA | 96.778 | 0 | 1391 | - |
| Chr7A_7343595 | TRINITY_DN8174_c0_g1_i7 | 45 | 97.342 | 2.55E-142 | 510 | Chaperone protein dnaJ 10 |
| Chr7A_7343595 | TRINITY_DN8174_c0_g1_i1 | 45 | 97.342 | 5.90E-142 | 510 | Chaperone protein dnaJ 10 |
| Chr7A_7343595 | TRINITY_DN8174_c0_g1_i2 | 45 | 97.342 | 6.22E-142 | 510 | Chaperone protein dnaJ 10 |
| Chr7A_7343595 | TRINITY_DN8174_c0_g1_i3 | 45 | 97.342 | 6.39E-142 | 510 | Chaperone protein dnaJ 10 |
| Chr8A_60112113 | TRINITY_DN4578_c0_g1_i7 | 35 | 99.281 | 4.95E-140 | 503 | RNA-binding protein with multiple splicing |
| Chr8A_60112113 | TRINITY_DN4578_c0_g1_i8 | 35 | 98.151 | 0 | 1122 | RNA-binding protein, mRNA-processing factor 2a |

| | | | | | | |
|---|---|---|---|---|---|---|
| Chr8A_60112113 | TRINITY_DN4578_c0_g1_i18 | 35 | 99.191 | 0 | 1783 | RNA-binding protein with multiple splicing |
| Chr8A_60112113 | TRINITY_DN4578_c0_g1_i19 | 35 | 99.191 | 0 | 1783 | RNA-binding protein with multiple splicing |
| Chr8A_60112113 | TRINITY_DN4578_c0_g1_i13 | 35 | 99.191 | 0 | 1783 | Cell wall integrity protein scw1 |
| Chr8A_60112113 | TRINITY_DN4578_c0_g1_i10 | 35 | 99.281 | 4.95E-140 | 503 | RNA-binding protein with multiple splicing |
| Chr8A_61045410 | TRINITY_DN5537_c0_g2_i1 | 9 | 96.183 | 0 | 641 | Adenylosuccinate lyase |
| Chr8A_61045410 | TRINITY_DN162679_c0_g1_i1 | NA | 94.416 | 3.50E-170 | 601 | - |
| Chr8A_61045410 | TRINITY_DN5537_c0_g3_i1 | NA | 96.614 | 0 | 832 | - |
| Chr8A_61807214 | TRINITY_DN6089_c0_g1_i5 | 15 | 99.631 | 2.01E-137 | 496 | - |
| Chr8A_61807214 | TRINITY_DN6089_c0_g1_i6 | 15 | 99.631 | 1.16E-137 | 496 | - |
| Chr8A_61807214 | TRINITY_DN6089_c0_g1_i7 | 15 | 99.631 | 1.11E-137 | 496 | - |
| tig00019374_12115 | TRINITY_DN9380_c0_g1_i4 | 17 | 93.516 | 0 | 137 | Xylan glycosyltransferase MUCI21 |
| tig00019634_1271 | TRINITY_DN19229_c2_g1_i1 | 1 | 92.529 | 5.28E-64 | 1566 | - |
| tig00019634_1272 | TRINITY_DN19229_c2_g1_i1 | 1 | 92.529 | 5.28E-64 | 1565 | - |
| tig00052406_31544 | TRINITY_DN12808_c0_g1_i2 | 1 | 97.081 | 0 | 706 | - |
| tig00052406_31581 | TRINITY_DN12808_c0_g1_i2 | 1 | 97.081 | 0 | 669 | - |
| tig00052406_31601 | TRINITY_DN12808_c0_g1_i2 | 1 | 97.081 | 0 | 649 | - |
| tig00080975_9059 | TRINITY_DN36687_c2_g1_i1 | 4 | 97.289 | 1.39E-158 | 1125 | - |
| tig00088430_738 | TRINITY_DN47576_c0_g1_i1 | NA | 97.81 | 9.04E-61 | 137 | - |

**Supplementary Figures**

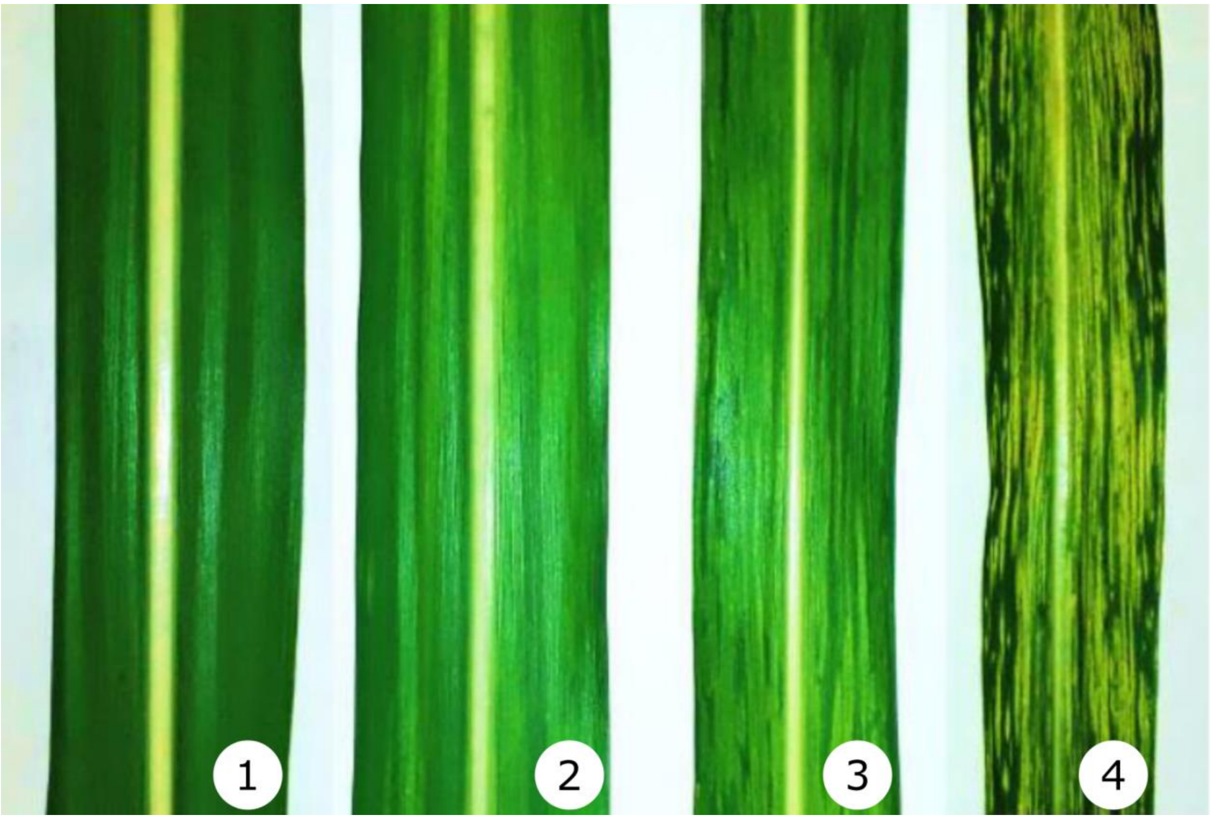


**Figure S1. Diagram of the scoring scale used for the assessment of sugarcane mosaic virus (SCMV) resistance based on the severity of mosaic symptoms.**

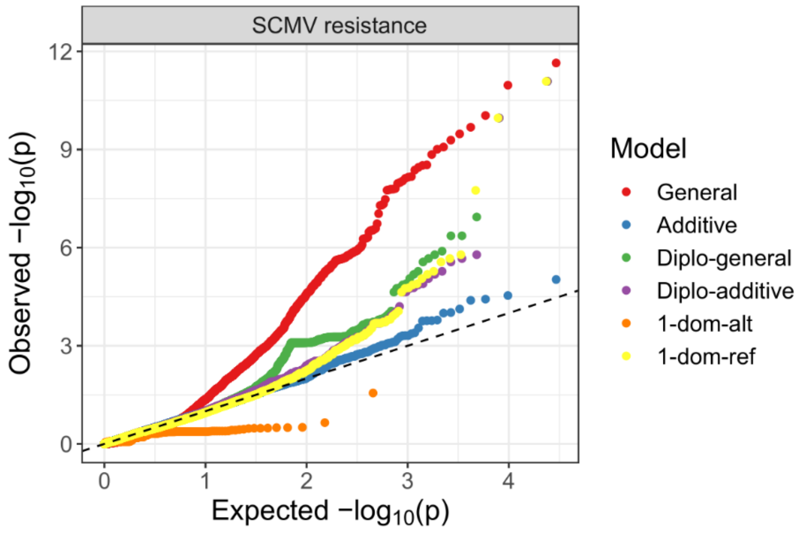


**Figure S2. Q-Q plots generated from the association analysis of sugarcane mosaic virus (SCMV) resistance.** Six different models were tested: general, additive, simplex dominant reference (1-dom-ref), simplex dominant alternative (1-dom-alt), diploidized general (diplo-general) and diploidized additive (diplo-additive) models.

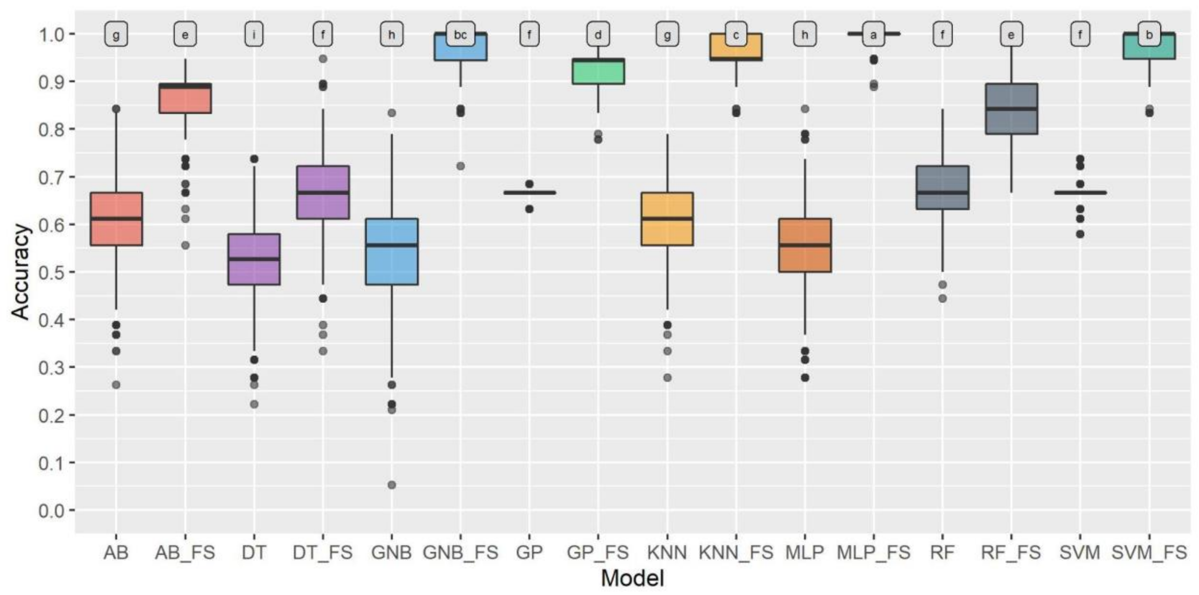


**Figure S3. Distribution of the accuracy of machine learning (ML) approaches employed to predict groups associated with sugarcane mosaic virus (SCMV) resistance when the full marker dataset and that obtained by feature selection (FS) were used.** The ML models tested were adaptive boosting (AB), decision tree (DT), Gaussian naive Bayes (GNB), Gaussian process (GP), K-nearest neighbor (KNN), multilayer perceptron neural network (MLP), random forest (RF) and support vector machine (SVM). The letters above boxplots indicate significantly different groups identified by ANOVA followed by Tukey’s test.

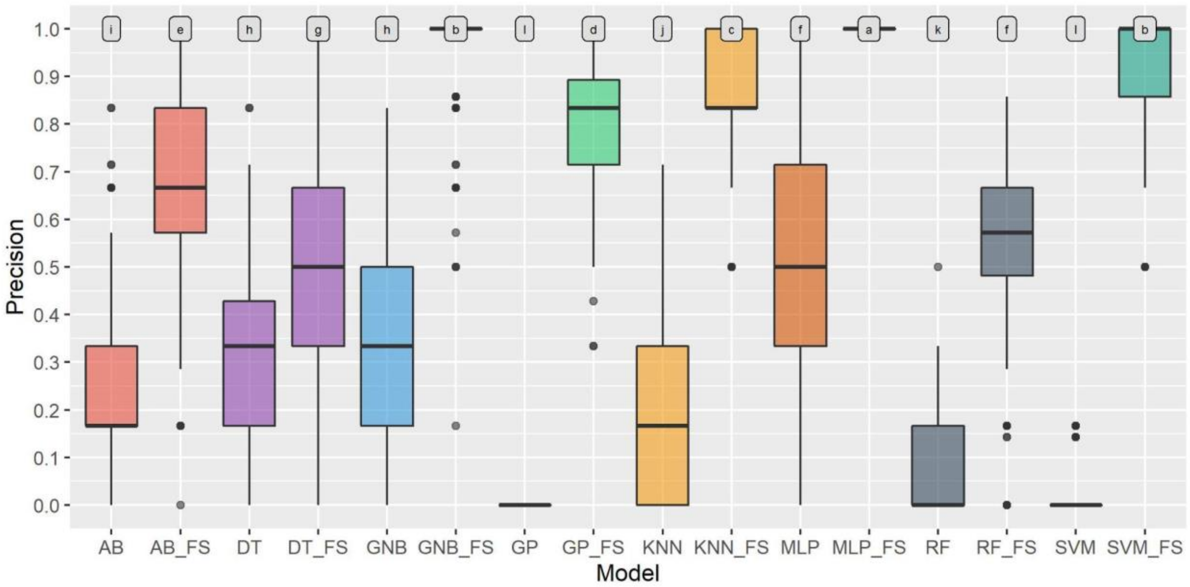


**Figure S4. Distribution of the precision of machine learning (ML) approaches employed to predict groups associated with sugarcane mosaic virus (SCMV) resistance when the full marker dataset and that obtained by feature selection (FS) were used.** The ML models tested were adaptive boosting (AB), decision tree (DT), Gaussian naive Bayes (GNB), Gaussian process (GP), K-nearest neighbor (KNN), multilayer perceptron neural network (MLP), random forest (RF) and support vector machine (SVM). The letters above boxplots indicate significantly different groups identified by ANOVA followed by Tukey's test.

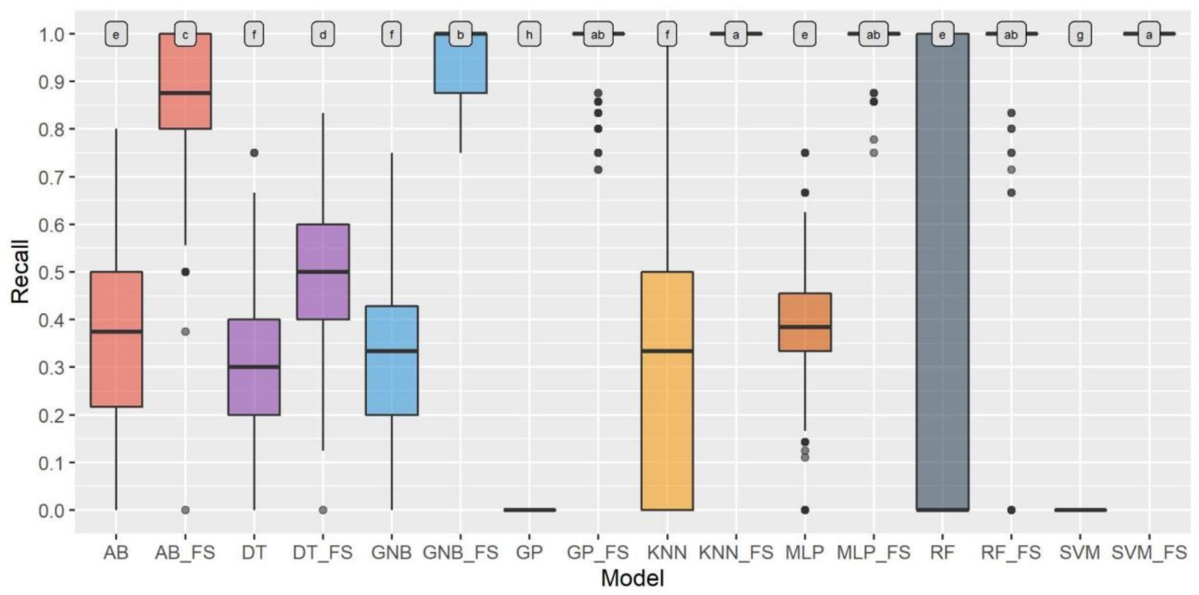


**Figure S5. Distribution of the recall of machine learning (ML) approaches employed to predict groups associated with sugarcane mosaic virus (SCMV) resistance when the full marker dataset and that obtained by feature selection (FS) were used.** The ML models tested were adaptive boosting (AB), decision tree (DT), Gaussian naive Bayes (GNB), Gaussian process (GP), K-nearest neighbor (KNN), multilayer perceptron neural network (MLP), random forest (RF) and support vector machine (SVM). The letters above boxplots indicate significantly different groups identified by ANOVA followed by Tukey's test.

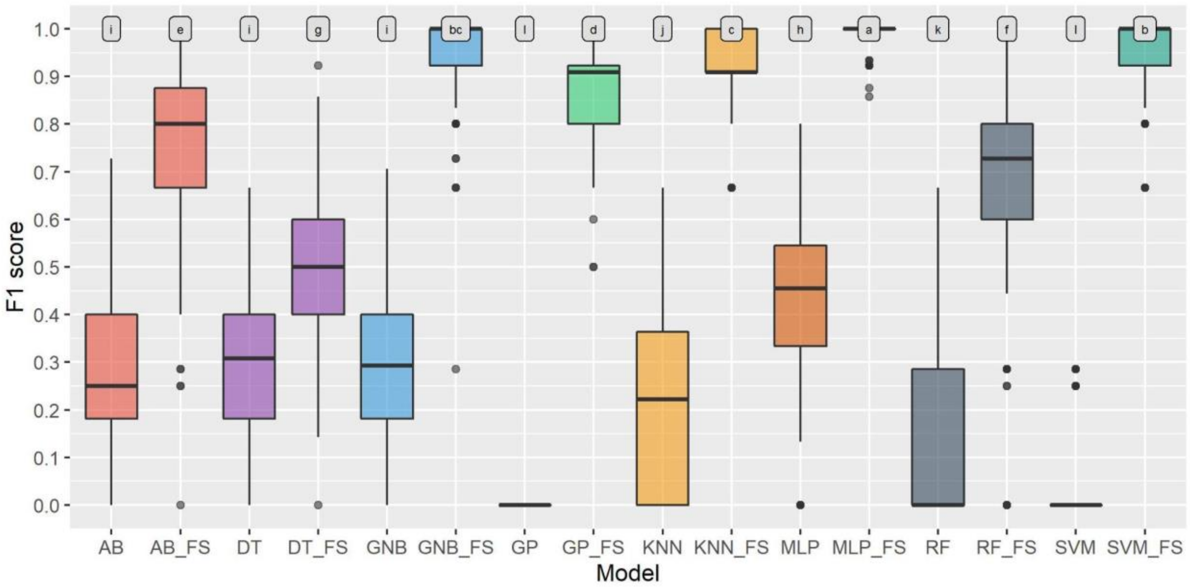


**Figure S6. Distribution of the F1 scores of machine learning (ML) approaches employed to predict groups associated with sugarcane mosaic virus (SCMV) resistance when the full marker dataset and that obtained by feature selection (FS) were used.** The ML models tested were adaptive boosting (AB), decision tree (DT), Gaussian naive Bayes (GNB), Gaussian process (GP), K-nearest neighbor (KNN), multilayer perceptron neural network (MLP), random forest (RF) and support vector machine (SVM). The letters above boxplots indicate significantly different groups identified by ANOVA followed by Tukey's test.

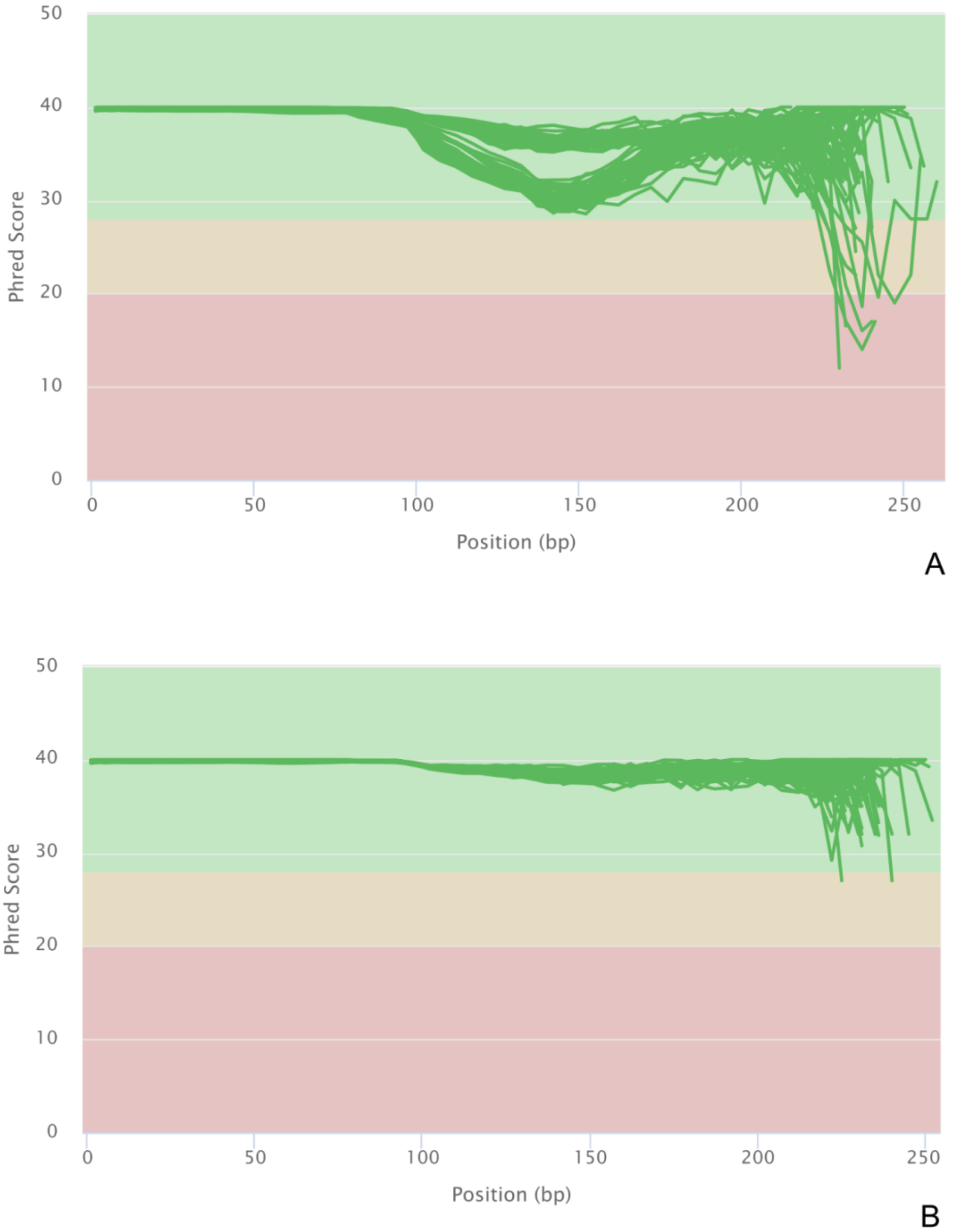


**Figure S7. Per-base sequence quality scores of raw (A) and trimmed (B) sequencing reads obtained via MonsterPlex technology.**

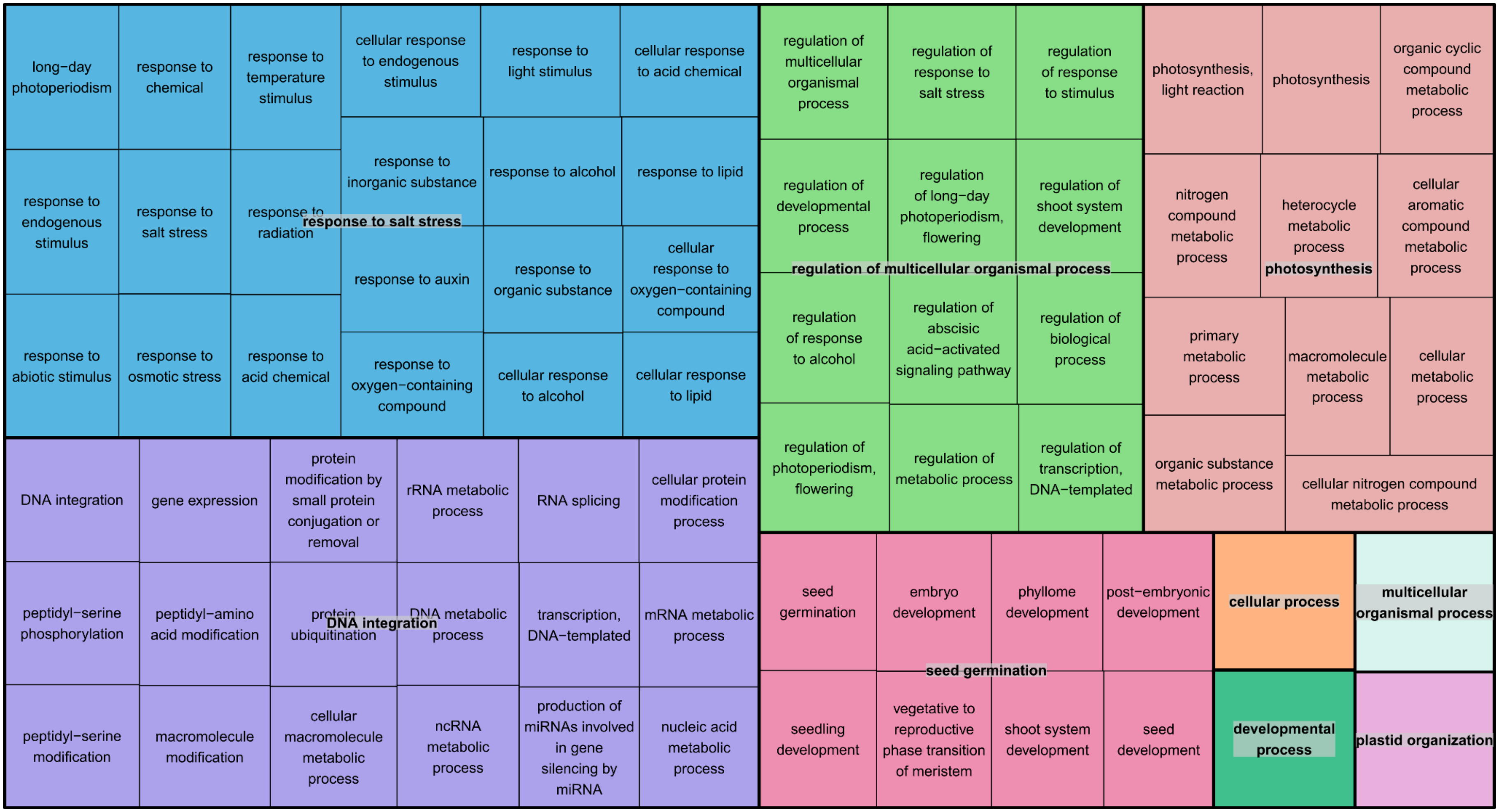


**Figure S8. TreeMap displaying Gene Ontology (GO) biological process enriched terms and categories obtained from all genes in the network modules that contained genes associated with sugarcane mosaic virus (SCMV) resistance.**
